# IL-2–induced Stat3 Signaling is Critical for Effector Treg Cell Programming

**DOI:** 10.1101/2023.09.26.559434

**Authors:** Emma C. Dean, Daniel F. Ditoro, Duy Pham, Min Gao, Carlene L. Zindl, Blake Frey, Stacey N. Harbour, David A. Figge, Aidan T. Miller, Caleb R. Glassman, K. Christopher Garcia, Robin D. Hatton, Casey T. Weaver

## Abstract

Maintenance of immune homeostasis to the intestinal mictrobiota is dependent on a population of effector regulatory T (eTreg) cells that develop from microbiota-reactive induced (i)Treg cells. A cardinal feature of eTreg cells is their production of IL-10, which plays a non-redundant role in immune tolerance of commensal microbes. Here, we identify an unexpected role for IL-2-induced Stat3 signaling to program iTreg cells for eTreg cell differentiation and *Il10* transcriptional competency. IL-2 proved to be both necessary and sufficient for eTreg cell development – contingent on Stat3 output of the IL-2 receptor coordinate with IL-2 signaling during early Treg cell commitment. Induction of iTreg cell programming in absence of IL-2-induced Stat3 signaling resulted in impaired eTreg cell differentiation and a failure to produce IL-10. An IL-2 mutein with reduced affinity for the IL-2Rγ (γ_c_) chain was found to have blunted IL-2R Stat3 output, resulting in a deficiency of *Il10* transcriptional programming that could not be fully rescued by Stat3 signaling subsequent to an initial window of iTreg cell differentiation. These findings expose a heretofore unappreciated role of IL-2 signaling that acts early to program subsequent production of IL-10 by developing eTreg cells, with broad implications for IL-2–based therapeutic interventions in immune-mediated diseases.

## Introduction

Pro-inflammatory effector (Teff) and anti-inflammatory regulatory CD4 T cells (Treg) exist in a dynamic equilibrium that prevents inappropriate inflammation while preserving the capacity to rapidly mount responses to invading pathogens. Post-thymic tolerance to self-antigens is maintained primarily by thymically derived (t)Treg cells, while tolerance to dietary and commensal microbial antigens is mediated by induced (i)Treg cells that develop from naïve precursors in peripheral lymphoid tissues (iTreg)^1–3^. Helios, encoded by *Ikzf2*, is a marker that distinguishes tTreg from iTreg cells^4^. Helios^+^ tTreg cells develop in the thymus from precursors with high affinity to self antigens and primarily seed lymphoid tissues, whereas Helios^-^ iTreg cells initially develop in peripheral lymphoid tissues, induced upon recognition of cognate foreign antigens to which tolerogenic responses are required^5^.

Different pathogens elicit distinct immune responses governed by phenotypically and functionally distinct subsets of effector CD4 T cells, each tailored to a specific inflammatory context. This functional specialization is reflected in the Treg cell compartment; Treg cells that express hallmark effector CD4 transcription factors (e.g., Rorgt, Gata3, Tbx21) are thought to be critical for controlling the corresponding effector subset (e.g., Th17, Th2, Th1)^6^. Maturation of both tTreg and iTreg cells into highly suppressive IL-10–producing “effector” Treg (eTreg) cells requires TCR ligation, cytokine stimulation, and induction of lineage-specific effector transcriptional networks^7^. Acquisition of additional anti-inflammatory properties by eTreg cells, particularly IL-10, is thought essential to enable eTreg cells to quell active T cell-driven inflammation.

At homeostasis, the intestines are home to the largest populations of Treg cells in the body, including Gata3^+^Helios^+^ tReg cells, and Rorγt^+^Helios^-^ and Rorγt^-^Helios^-^ iTreg cells^8–13^. Intestinal Helios^+^ tTreg cells are important in tissue repair and maintenance of the intestinal epithelium^9,10^. In contrast, Helios^-^ iTreg cells appear to be tasked with reinforcing anti-inflammatory responses to ingested or commensal microbe-derived antigens. The Rorγt^+^Helios^-^ subset of iTreg cells are a highly suppressive, terminally differentiated subset of eTreg cells that is required for inhibiting Th17-mediated inflammation^13–15^. These cells – and IL-10^+^ Treg cells in general – are found in highest frequencies in the large intestine (LI), which contains the greatest density and diversity of commensal microbes^16^. These cells are strictly dependent on expression of IL-10 to maintain homeostasis; loss of this iTreg population or its expression of IL-10 causes lethal colitis^17,18^. Yet our understanding of the signals required for the development and mantenance of this critical Treg population remains incomplete.

Expression of Rorγt is regulated by signal transducer and activator of transcription 3 (Stat3) and is required for the suppressive function of Rorγt^+^Helios^-^ iTreg cells^19^. The balance in favor of Stat3 or Stat5 activation during early T cell differentiation skews toward Th17 or Treg cells, respectively; Stat3 is required for expression of Rorgt in both Teff and Treg cells, yet Stat3 is antagonistic to developing Treg cells^20–23^. The contrasting actions of Stat3 in Treg cell biology, and the ligands driving its activation, are poorly understood. Stat5, in contrast, promotes Treg development^24,25^. Interleukin-2 (IL-2) is the primary mediator of Stat5 signaling in Treg cells and is required for their development and maintenance^26,27^. Yet IL-2 can also act on Teff cells to promote, rather than repress, T cell-driven inflammation^28^. Accordingly, efforts to target IL-2 specifically to Treg cells have gained interest. IL-2 complexes and synthetic IL-2 muteins that, through altered interactions with IL-2 receptor (IL-2R) components, specifically signal on Treg cells are in development for therapeutic applications in immune-mediated disease^29–31^. However, efforts to disentangle the role of IL-2 in the differentiation and function of Treg subsets have been complicated by its broad role in Treg cell biology; the specific role of IL-2 signaling in the development of tissue iTreg cells – and eTreg cells in particular – remains to be defined.

Here, through exploration of the transcriptional programs that distinguish large intestinal Treg cells that do or do not express *Il10* transcripts, we have exposed signaling pathways required for the induction of IL-10^+^ intestinal effector iTreg cells. We find that Stat3 signaling is specifically enriched in IL-10–competent iTreg cells in the LI, and that Stat3 signaling is required to support the development of these intestinal eTreg cells. Further, we identify IL-2 as the major source of Stat3 activation driving differentiation of this population. Our findings uncover an unappreciated role of IL-2 signaling that acts early to program subsequent production of IL-10 by effector iTreg cells, with broad implications for IL-2–based therapeutic interventions in immune-mediated diseases.

## Results

### Colonic IL-10 iTreg cells are phenotypically distinct from thymically derived populations

The large intestine has the highest frequency of IL-10 producing Treg cells in the body^16^, and they are heterogenous^32,33^. To define subset specificity of IL-10 expression and determine whether IL-10 competent cells might rely on different signals for their development, we performed single-cell RNA-sequencing on total colonic TCRβ^+^CD4^+^ T cells from C57BL/6 mice colonized with *Helicobacter typhlonius*, which has been shown to increase colonic eTreg populations^14^ (**Fig. 1a, Fig. S1a**). Experiments performed using *H. typhlonius*-negative animals found similar populations albeit in smaller numbers (data not shown).

**Fig. 1.**
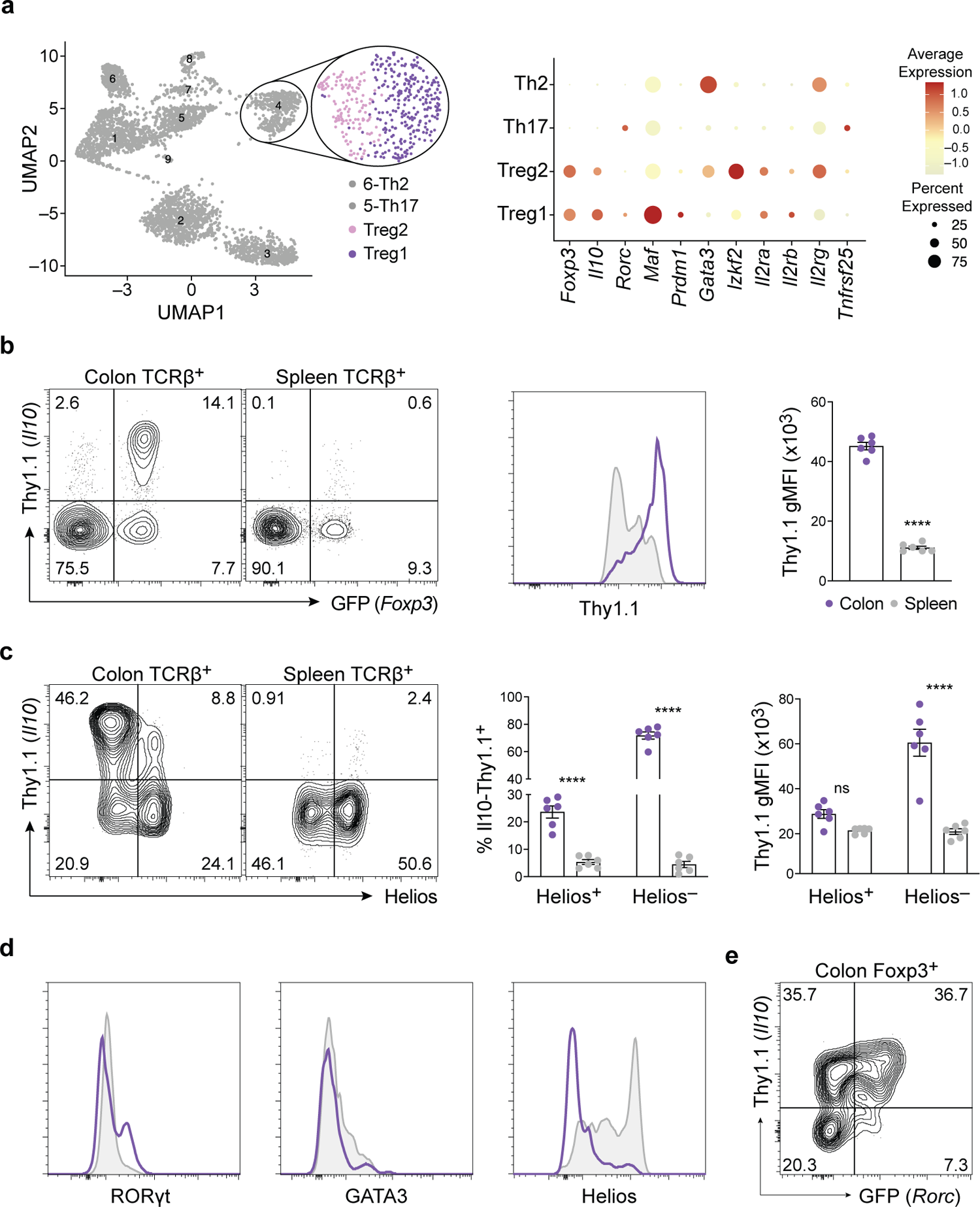
Helios^-^ iTreg cells comprise the IL-10 competent pool in the colon. **a,** Single cell RNA-seq was performed on live CD4^+^TCRβ^+^ T cells isolated from the colonic lamina propria of naïve C57BL/6 mice. Cluster analysis (left) depicts all cells sequenced. Dot plot (right) depicts average expression of indicated genes in select clusters. Data are representative of two independent experiments. **b-d,** Lymphocytes were isolated from the colonic lamina propria (purple) and spleens (grey) of 10BiT.*Foxp3*GFP mice and assessed by flow cytometry. Flow cytometric profiles (left) depict cell number controlled concatenated averages of biological replicates (gated on CD4^+^TCRβ^+^ cells). Experiments performed twice. **b**, Expression of Thy1.1 and *Foxp3*.GFP (left). Histogram depicting Thy1.1 expression (gated on CD4^+^TCRβ^+^*Foxp3*GFP^+^ cells) (middle). Quantification depicts Thy1.1 expression geometric mean fluorescence intensity (gMFI) (right) (n=6 for both colon and spleen, mean ± SEM; ****P<0.0001). **c,** Expression of Thy1.1 and Helios (left). Quantifications depict percent of tTreg (CD4^+^TCRβ^+^Foxp3^+^Helios^+^) and iTreg (CD4^+^TCRβ^+^*Foxp3*GFP^+^Helios^-^) expressing Thy1.1^+^ (middle) and Thy1.1 gMFI among tTreg versus iTreg cells (n=6 for both colon and spleen, mean ± SEM; ns, nonsignificant, ****P<0.0001). **d,** Expression of Rorγt, Gata3, and Helios in Thy1.1^+^ Treg cells (gated on CD4^+^TCRβ^+^*Foxp3*GFP^+^Thy1.1^+^). Histograms are representative of three independent experiments. **e,** Lymphocytes were isolated from the colonic lamina propria of 10BiT.*Rorc.*eGFP mice and assessed by flow cytometry. Data depict cell number controlled concatenated averages of biological replicates. Experiment performed twice. Statistical differences were tested using unpaired Student’s t-test (two-tailed) and two-way ANOVA.

The transcriptomic data identified cells of the major T helper subsets, including Th1, Th2, and Th17 cells, as well as two distinct subpopulations of Treg cells: Treg1 and Treg2 (**Fig. 1a, Fig. S1a**). The Treg2 cluster was enriched for genes associated with tTreg cells, including *Gata3*, *Ikzf2* (Helios-encoding) and components of the IL-2 receptor complex, particularly the gene encoding the γ_c_ chain. Conversely, the Treg1 cluster was enriched for genes associated with effector iTreg cells, including *Il10, Maf, Prdm1,* and *Rorc*^13–15,34^ (**Fig. 1a**). The majority of *Il10* expressing cells did not express *Ikzf2*, though a small sub-cluster of cells expressed both *Il10* and *Ikzf2* (**Fig. S1b**). These data indicated that there were two dominant subsets of Treg cells in the colon that were divergent based on IL-10 expression. These two populations also appeared to diverge based on IL-2 signaling components, with the Treg2 subset particularly enriched for transcripts that encode the γ_c_ chain. Our findings are consistent with recent studies that identified a suppressive colonic Treg cell population characterized by expression of *Il10, Gzmb, Lag3,* and *Cxcr3*, which resembled Rorγt-expressing iTreg cells^32^.

To validate these data, we interrogated IL-10 expression in colonic Treg cells using dual–reporter 10BiT.*Foxp3*GFP mice, which bear transgenes for both a *Foxp3*GFP reporter^35^ and a surface Thy1.1 reporter that faithfully reports reports active (Thy1.1^bright^), inactive (Thy1.1^intermediate/low^) or no (Thy1.1^negative^) expression of *Il10*^16^. As Treg cells in peripheral lymphoid tissues are mostly tTreg cells, comparison of colonic Treg cells to splenic Treg cells allowed us to further determine whether IL-10 was a suitable marker to segregate colonic Treg cell subsets by cell of origin^36^. FACS analysis showed a large population of Thy1.1^bright^ cells in the colon not found in the spleen; consequently, the average Thy1.1 gMFI in the colonic Treg cells was significantly higher than in splenic Treg cells (**Fig. 1b**).

To determine whether iTregs comprised the Thy1.1^bright^ group, we bifurcated the cells based on Helios expression and analyzed Thy1.1 gMFI. Splenic Treg cells were predominantly Helios^+^ and the limited Thy1.1 expression observed (prdominantly Thy1.1^intermediate/low^) was largely restricted to these Helios^+^ cells (**Fig. 1c, S1c**), which were characterized by elevated Gata3 levels (**Fig. 1d**). In contrast, significantly lower frequencies of Helios^+^ Treg cells were found in colon, and nearly all Thy1.1^bright^ cells were Helios^-^ iTreg cells (**Fig. 1c, S1c**); essentially no Helios^+^ Treg cells in colon were were Thy1.1^bright^. Rorγt expression was highly enriched in colonic Helios^-^IL-10^+^ cells (**Fig. 1d**); about half of colonic Thy1.1^+^Helios^-^ Treg cells expressed Rorγt (**Fig. 1e**). This specific subset of Treg cells is induced in response to the microbiota and is important in maintaining the anti-inflammatory tone in the intestine^13–15^. Similar results were found using SPF mice lacking the *H. typhlonius* pathobiont (not shown). Taken together, these data indicates that: IL-10 suitably bifurcates the two major populations of Treg cells in the colon; IL-10 competent Treg cells in the colon produce higher amounts of IL-10 compared to those in the spleen; and IL-10–competent colonic Treg cells are prinicpally derived from microbiota-reactive peripherally induced iTreg cells.

### Stat3 signaling drives IL-10 production in colonic iTreg cells

Prior studies have compared the transcriptomes of lymphoid and colonic Foxp3^+^ cells^8,32^ but signaling pathways that may be differentially utilized by IL-10 competent cells in each tissue have not been reported. The transcriptional and phenotypic differences observed between splenic and colonic IL-10 competent cells suggested there were signaling pathways uniquely enriched in iTreg versus tTreg cells. To explore this further, bulk RNA-sequencing of colonic and splenic Thy1.1^+^ and Thy1.1^-^ Treg cells was performed. Thy1.1^+^ colonic Treg cells mirrored the Treg1 subgroup identified by single cell RNA-sequencing; the IL-10 competent cells expressed higher levels of *Prdm1* and *Rorc*, and lower levels of *Gata3*, *Ikzf2, Il2ra, Il2rb,* and *Il2rg* (**Fig. 2a and Fig S2**). These data indicated that Thy1.1 expression by colonic Treg cells distinguishes the two major populations identified by single cell transcriptomic analyses, and that the IL-10 competent cells were mostly iTreg cells. In contrast, while splenic Thy1.l^+^ cells exhibited modest increases in *Prdm1* and *Rorc* expression, they were characterized by substantially elevated *Gata3*; further, there was no difference in the expression of *Ikzf2* between Thy1.1^+^ and Thy1.1^-^ cells. In line with flow cytometric data, these findings confirm that splenic IL-10–competent cells were predominantly derived from tTreg cells.

**Fig. 2.**
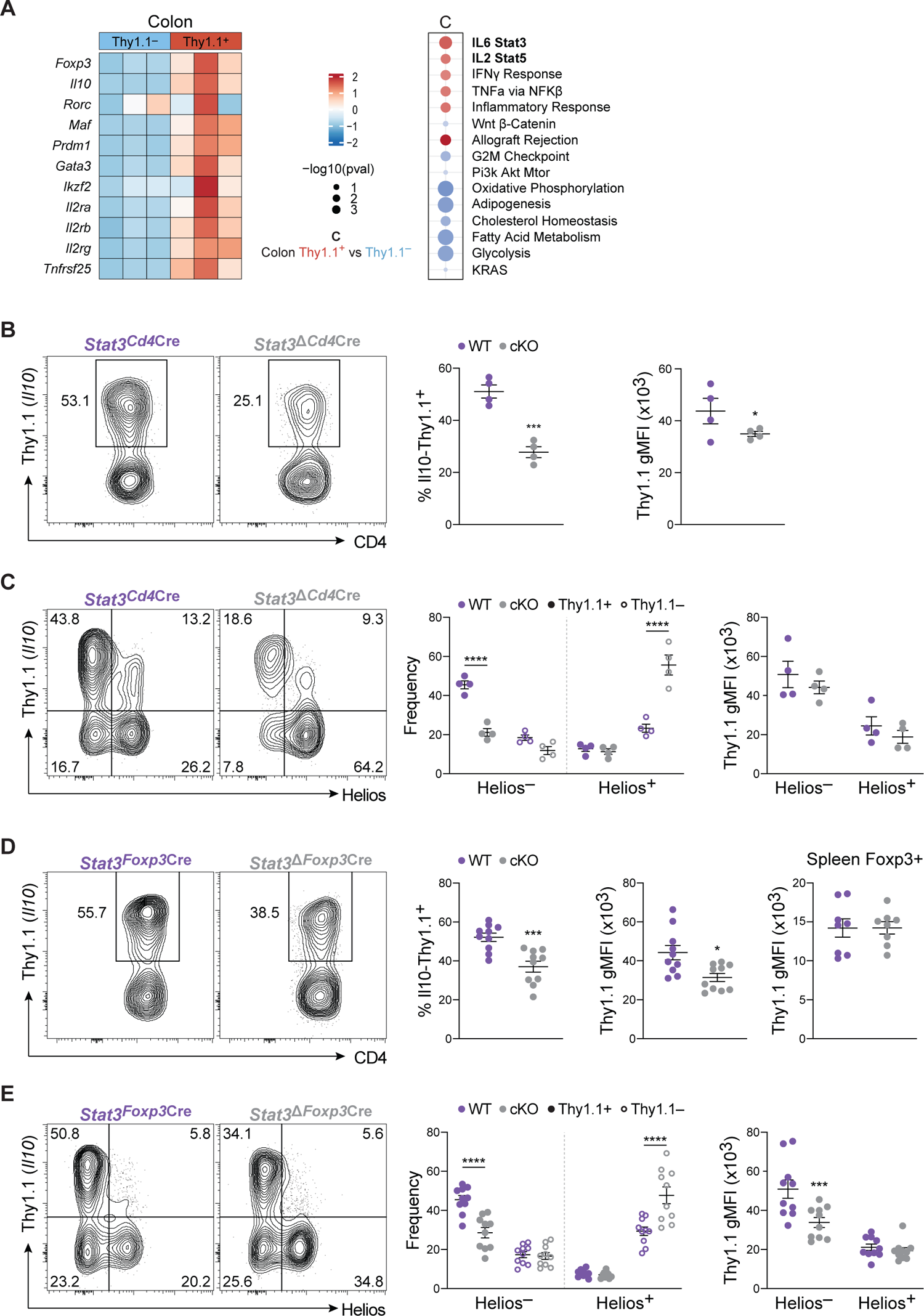
IL-10 competent colonic iTreg cells are enriched for Stat3 signaling. **a**, RNA sequencing was performed on CD4^+^TCRβ^+^*Foxp3-*GFP^+^Thy1.1^-^ (light blue) and CD4^+^TCRβ^+^*Foxp3-*GFP^+^Thy1.1^+^ (red) Treg cells sorted from colonic lamina propria of 10BiT.*Foxp3-GFP* mice. Heatmap (left) depicts relative expression (z-score) of select genes between Thy1.1^-^ and Thy1.1^+^ Treg cells from the colon (n=3). Gene set enrichment analysis (GSEA, right) depicts enriched hallmark gene sets in Thy1.1^+^ versus Thy1.1^-^ Treg cells from the colon. **b-c,** Treg cells were isolated from the colonic lamina propria of gender matched *Stat3*^CD4Cre^ (WT, purple) and *Stat3*^ΔCD4Cre^ (cKO, grey) littermate mice and assessed by flow cytometry. Cytometric profiles depict cell number controlled concatenated averages of biological replicates. **b,** Thy1.1 expression in Treg cells (left, gated on CD4^+^TCRβ^+^Foxp3^+^). Quantification showing percent Thy1.1^+^ (middle) and Thy1.1^+^ gMFI (right) in Treg cells between WT and cKO (n=4, mean ± SEM; *P<0.05, ***P<0.001). **c**, Thy1.1 and Helios expression in colonic Treg cells (left, gated on CD4^+^TCRβ^+^Foxp3^+^). Quantification showing Thy1.1 gMFI (middle) and frequency (right) in tTreg and iTreg cells (n=4, mean ± SEM; ****P<0.0001). **d-e,** Treg cells isolated from the colonic lamina propria or spleen of geneder matched *Stat3*^Foxp3Cre^ (WT, purple) and *Stat3*^ΔFoxp3Cre^ (cKO, grey) littermate mice and assessed by flow cytometry. Cytometric profiles depict cell number controlled concatenated averages of biological replicates. **d,** Thy1.1 expression in Treg cells (left, gated on CD4^+^TCRβ^+^*Foxp3*YFP^+^). Quantifications showing percent Thy1.1^+^ (first) and Thy1.1^+^ gMFI (second) in colon Treg cells and Thy1.1^+^ gMFI (third) in spleen Treg cells (n=10 for colon and spleen, mean ± SEM; *P<0.05, ***P<0.001). **e,** Thy1.1 versus Helios expression in colonic Treg cells (left, gated on CD4^+^TCRβ^+^*Foxp3*YFP^+^). Quantifications showing Thy1.1 gMFI (middle) and frequency (right) in colonic tTreg and iTreg cells (n=10, mean ± SEM; ***P<0.001, ****P<0.0001). Statistical differences were tested using unpaired Student’s t-test (two-tailed) and two-way ANOVA.

To explore signaling mechanisms unique to colonic IL-10–competent cells, we performed gene set enrichment analysis (GSEA) on genes differentially expressed (DEGs) between Thy1.1^+^ and Thy1.1^-^ Treg cells in the colon and spleen (**Fig. 2a, S2b**). Both IL-2 Stat5 and IL-6 Stat3 pathways were enriched in IL-10–competent colonic Treg cells, indicating that iTreg cells may rely on these cytokine signals for their development and/or function. It has been well established that Stat5 signaling is critical for Treg cell development, maintenance, and function, while Stat3 has been shown to be generally antagonistic to the Treg cell program^20–23,37^. However, studies have also shown that Stat3 expression plays a critical role in Treg control of Th17–mediated inflammation in the intestine^11,19^. From these data and our own, we hypothesized that Stat3 activation mediated by cytokine signaling was critical for IL-10 competency in colonic iTreg cells.

To examine the effects of Stat3 deficiency *in vivo* we induced conditional deletion of *Stat3* by crossing mice carrying a floxed *Stat3* allele (*Stat3^fl/fl^*)^38^ to *CD4*Cre mice that also carried the 10BiT reporter (“*Stat3*^Δ*CD4*Cre^”) (**Fig. 2b-c**). Compared to wild type littermate controls (*Stat3^CD4^*^Cre^), *Stat3*^Δ*CD4*Cre^ mice had a significant decrease in the frequency of colonic Thy1.1^+^Foxp3^+^ Treg cells due to a deficit in the Thy1.1^+^Helios^-^ subgroup. Interestingly, there was a concomitant increase in the proportion of Thy1.1^-^Helios^+^ cells (**Fig. 2c**). Thy1.1gMFI was reduced in both Helios^-^ and Helios^+^ subsets (**Fig. 2b-c**). Stat3 controls Rorγt expression in Th17 cells^39^; accordingly, we found essentially no Rorγt^+^ Treg cells and a reduction in Th17 cells in the *Stat3*^Δ*CD4*Cre^ animals (**Fig. S2c**).

Because Stat3 is critical in the development and function of Th17 cells and Rorγt expressing Tregs are critical in controlling Th17-mediated inflammation, there could have been cell–extrinsic factors affecting these results. We therefore sought to examine the role of Stat3 exclusively in Treg cells using *Stat3*^Δ*Foxp3*Cre^ mice. These mice harbor the 10BiT reporter with *Stat3* conditionally deleted in *Foxp3*YFPCre–expressing cells. *Stat3*^Δ*Foxp3*Cre^ mice showed similar changes in Treg cell populations as *Stat3*^Δ*CD4*Cre^ mice. However, the decrease in Thy1.1 gMFI was restricted to colonic Helios^-^ cells (**Fig. 2d-e**). Accordingly, we saw no changes in Thy1.1gMFI in splenic Treg cells, which are mostly Helios^+^ tTreg cells (**Fig. 2d**). There was also an increase in IL-17A–producing cells (**Fig. S2d**), is line with published data that *Stat3*^Δ*Foxp3*Cre^ mice develop spontaneous inflammation^19^. Taken together, these data indicated that *Stat3* deficiency significantly reduced IL-10 production by colonic iTreg cells via a cell-intrinsic mechanism, indicating an important contribution of this signaling pathway to this subpopulation.

### iTreg cells require early Stat3 signaling to program IL-10 competency

A major barrier to studying eTreg cells has been the lack of *in vitro* systems for generation of sufficient numbers of cells with phenotypic and functional fidelity to eTreg cells that develop *in vivo.* Building on a recent report showing that vitamin C acts with Tet2 to stabilize *Foxp3* expression *in vitro*^40^ through sustained demethylation of CNS2 within *Foxp3*^13,41^, we found that addition of vitamin C during iTreg polarization from naive CD4 T cell precursors enabled us to generate a high frequency of IL-10–competent Treg cells *in vitro*. This required a two–stage culture process that included both initial polarization and restimulation phases (**Fig. 3a**). The iTreg cells generated using this protocol were able to cure Th17 transfer colitis^42^ (**Fig. 3b**), harbored a gene expression signature consistent with eTreg fate determination, and were enriched for Stat3 signaling in the IL-10 competent subset (**Fig. 3c**). Thus, iTreg cells produced via this *in vitro* system recapitulated critical features of peripherally induced Treg cells *in vivo*, including their high expression of IL-10, and affirmed that Stat3 signaling might be important eTreg cell development.

**Fig. 3.**
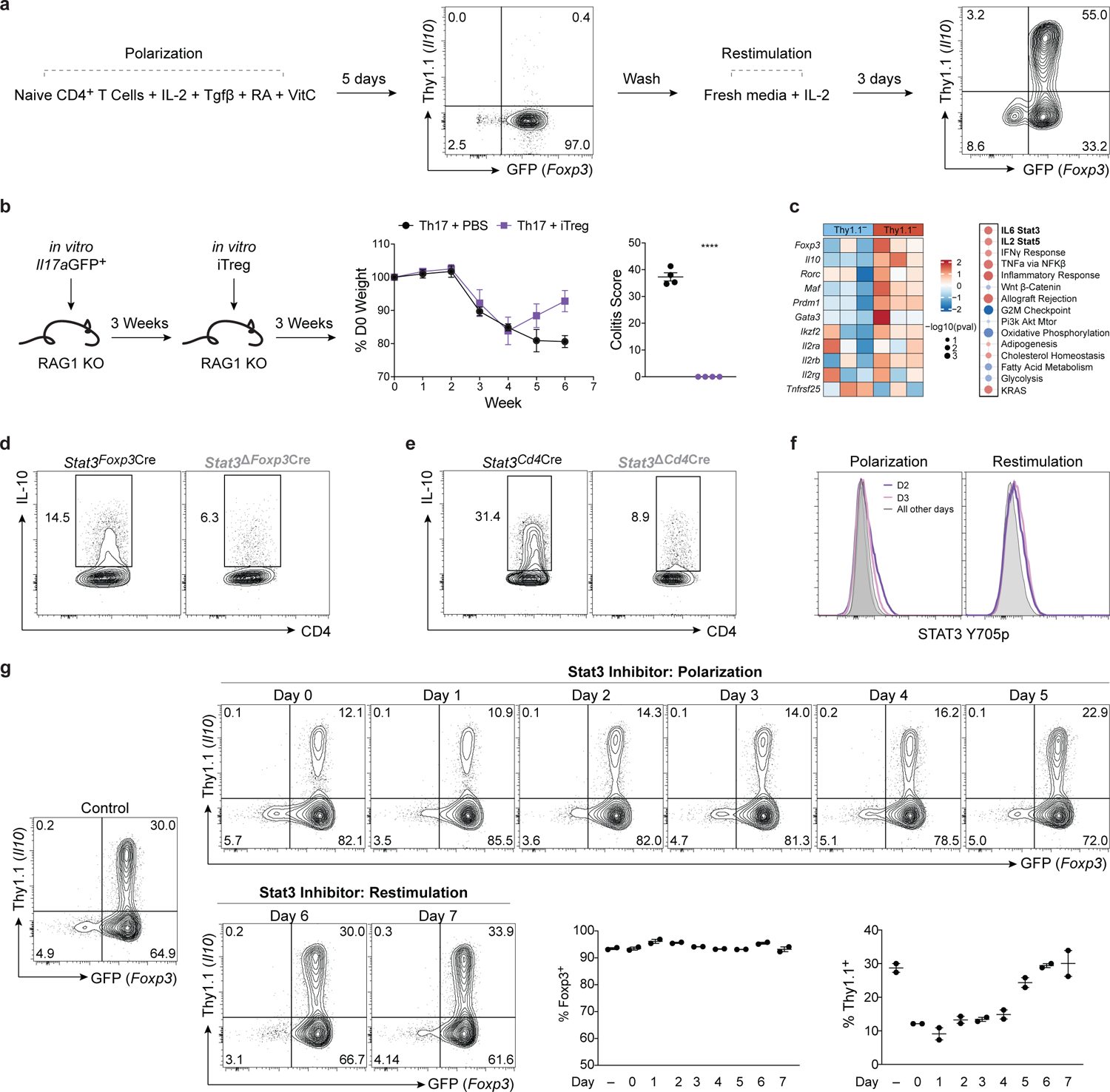
IL-10 competent *in vitro* iTreg cells rely on Stat3 signaling early in their development. **a**, *In vitro* derived iTreg cell polarization schematic, using 50U/mL IL-2. Flow cytometric profiles (middle, right) show Thy1.1 versus *Foxp3*GFP expression in CD4+ T cells at days 5 and 8, respectively. **b,** Naïve CD4 T cells from *Il17a*.GFP mice were stimulated *in-vitro* in Th17 conditions. Purified *Il17a*.GFP+ Th17 cells were transferred into *Rag1* KO mice (CD45.2) to induce colitis. *in vitro* derived d5 iTreg cells (CD45.1) were transferred after 3 weeks. Mice were weighed weekly (middle). After 6 weeks, animals were sacrificed and tissue was scored for histpathology (right) (n=4, mean ± SEM; ****P<0.0001). **c,** RNA-sequencing was performed on *in vitro* derived Thy1.1^+^ and Thy1.1^-^ *Foxp3*GFP^+^ Treg cells after 8 days of stimulation as in Fig. 3a. Heatmaps (left) showing relative expression (z-score) of target genes between Thy1.1^-^ and Thy1.1^+^ Treg cells (n=3). Gene set enrichment analysis (GSEA, right) showing enriched hallmark gene sets in Thy1.1^+^ versus Thy1.1^-^ Treg cells. -log10(pval) of 1 represents a p value < 0.1 (not significant), -log10(pval) of 2 represents a p value < 0.01, and -log10(pval) represents a p value <0.001. Data represent the combined analysis of three biologically independent samples. **d-e,** Naïve CD4 T cells were isolated from gender matched littermate *Stat3*^CD4Cre^ (black) and *Stat3*^ΔCD4Cre^ (grey) (**d**) or *Stat3*^Foxp3Cre^ (black) and *Stat3*^ΔFoxp3Cre^ (grey) (**e**) mice and polarized in iTreg conditions for 8 days as in Fig. 3a then assessed by flow cytometry for expression of *Il10*. Flow cytometric profiles depict cell number controlled concatenated averages of technical replicates. Experiments performed 3 times. **f,** Histograms showing kinetic staining of Stat3 Y705p during iTreg polarization (left) and re-stimulation (right). Flow cytometric profiles depict cell number controlled concatenated averages of technical replicates. Experiment performed 3 times. **g,** Naïve CD4 T cells were isolated from 10BiT.*Foxp3*GFP mice and stimulated in iTreg conditions as in Fig. 3a. but 42µM SD-36 (Stat3 inhibitor) or DMSO (control, left) was added on each day to independent wells (top row identifies days in the polarization phase, bottom row shows days in the re-stimulation phase). Flow cytometric profiles depict Thy1.1 and *Foxp3*-GFP expression on day 8 of stimulation. Quantifications depict frequency of *Foxp3*GFP^+^ (middle) and Thy1.1^+^ (right) cells on day 8 (n=2). Statistical differences were tested using unpaired Student’s t-test (two-tailed). Experiment performed twice.

To examine the role of Stat3 in the development of these induced eTreg cells, we polarized naïve CD4 T cells from *Stat3*^Δ*Foxp3*Cre^ mice and littermate controls. Loss of Stat3 led to ∼50% reduction in IL-10 expression (**Fig. 3d**). The same experiment performed using cells from *Stat3*^Δ*CD4*Cre^ mice, however, led to a more substantial defect in IL-10 production (**Fig. 3e**). Stat3 is deleted in T cells of CD4Cre mice during thymic development, whereas Foxp3Cre mice do not induce loss of Stat3 until Foxp3 is expressed during differentiation of iTreg cells from naïve CD4 T cell precursors. The discrepancy in ablation of IL-10 expression observed suggested that Stat3 signaling may be required early in effector iTreg cell development to program these cells for IL-10 competency.

To explore this, we measured the kinetics of Stat3 activation across the polarization and re-stimulation phases of the iTreg cell culture system. We found that Stat3 phosphorylization peaked approximately 48 hours after TCR stimulation in both phases (**Fig. 3f**). These data indicated that Stat3 signaling may be important in either priming these cells for an effector phenotype, modulating IL-10 expression after re-stimulation, or both. We then polarized iTreg cells *in vitro* and spiked in the Stat3 inhibitor SD-36 at each day of the polarization (**Fig. 3g**). SD-36 is a highly specific Stat3 inhibitor that prevents homodimerization and translocation to the nucleus^43^. We utilized a dose that significantly reduced Stat3 activation without inducing unwanted off-target and toxic effects (**Fig. S3a**). Interestingly, we found that IL-10 competency was only affected when the inhibitor was added within the polarization phase (first 5 days) (**Fig. 3g**); we saw no effect on Treg IL-10 competency when the inhibitor was added during the re-stimulation phase, despite a clear reduction in Stat3 phosphorylation (**Fig. S3b**). These data indicate that Stat3 signaling early during iTreg cell development is critical in programming subsequent IL-10 competency of iTreg cells.

### IL-2–induced Stat3 activation drives effector iTreg cell development *in vitro*

Stat activation in T cells is downstream of cytokine receptor signaling^44^; as such, pStat levels are a direct readout of the cytokine milieu in which T cells are stimulated and induced. Alhtough IL-2 is established as a requisite cytokine for the development and maintenance of Treg cells, this is typically attributed to its activation of Stat5. However, our RNA-sequencing, *in vivo*, and *in vitro* data implicated Stat3 signaling as critical for IL-10 competency in effector iTreg cells. We therefore sought to determine which cytokines were responsible for Stat3 activation in colonic eTreg cells. Because we observed Stat3 activation in the context of high levels of Stat5 activation, and both of our transcriptomic analyses identified IL-2R signaling as differentially utilized by iTreg and tTreg cells, we reasoned that IL-2, a cytokine capable of activating both Stat5 and Stat3^45^, may be important in this context.

Because the only exogenous Stat-activating cytokine in our *in vitro* iTreg system was IL-2, we assessed IL-2–induced activation of Stat3 over a titrated dose range during the initial eTreg polarization. IL-2 activated both Stat3 and Stat5 in a dose-dependent manner (**Fig. 4a**). Importantly, IL-10 competency at day 8 was dependent on the concentration of IL-2 used during the initial polarization and not during restimulation (**Fig. 4a and S4A**). These data indicated that, like Stat3, early exposure of Treg cells to IL-2 signaling dictated their effector capacity upon re-stimulation.

**Fig. 4.**
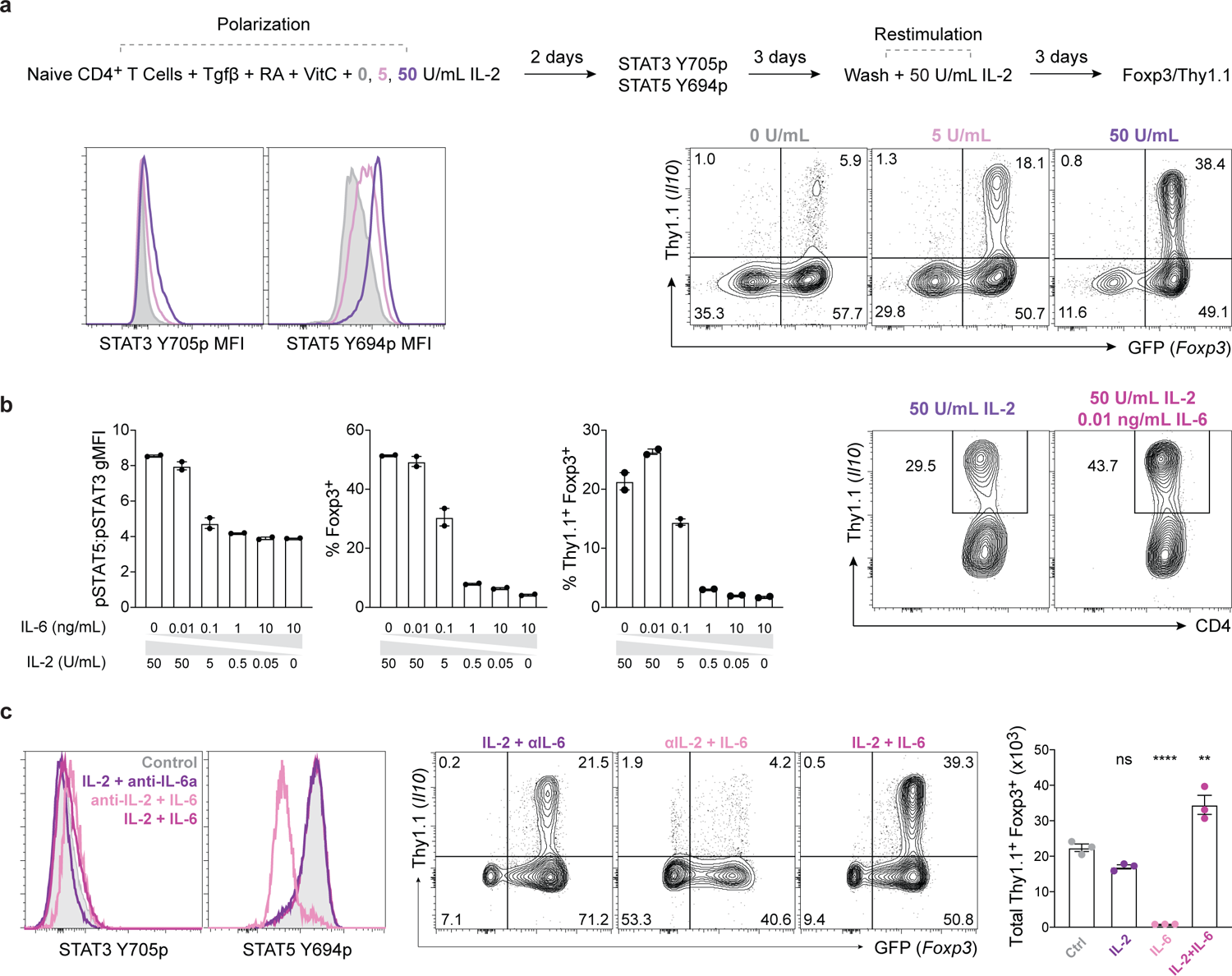
IL-2-induced Stat3 activation is sufficient for robust eTreg cell development. **a**, Naïve CD4 T cells were isolated from 10BiT.*Foxp3*GFP mice and polarized in vitro in iTreg conditions using 0 (grey), 5 (pink) or 50 (purple) U/mL IL-2 during the induction phase (left) and 50 U/mL during re-stimulation. Histograms (left) showing Stat3 Y705p and Stat5 Y694p at day 2 for each dose of IL-2 (gated on live CD4^+^*Foxp3*GFP^+^). Flow cytometric profiles showing Thy1.1 versus *Foxp3*GFP expression on day 8 for each dose of IL-2 (bottom right) (gated on live CD4^+^) depict cell number controlled concatenated averages of technical repeats. Experiment performed 3 times. **b,** Naïve CD4 T cells were isolated from 10BiT.*Foxp3*GFP mice and polarized in vitro in iTreg conditions using the indicated concentrations of IL-2 and IL-6 during the induction phase and 50 U/mL during re-stimulation. Quantifications show the ratio of Stat3 Y705p to Stat5 Y694p in *Foxp3*GFP^+^ Treg cells on day 2 of polarization (left), the frequency of *Foxp3*GFP^+^ cells on day 5 (middle), and the frequency of Thy1.1^+^*Foxp3*GFP^+^ cells on day 8 (right). Representative flow cytometric profiles showing Thy1.1 versus CD4 expression (gated on *Foxp3*GFP^+^) at day 8 (far right). **c,** Naïve CD4 T cells were isolated from 10BiT.*Foxp3*GFP mice and polarized in vitro in iTreg conditions with 50U/mL IL-2 (Control, grey), 50U/mL IL-2 + 10µg/mL αIL-6Rα (IL-2 + anti-IL-6a, purple), 10µg/mL αCD25 nd 0.1ng/mL IL-6 (anti-IL-2 + IL-6, pink), or 50U/mL IL-2 + 0.1ng/mL IL-6 (IL-2 +IL-6, magenta). Histograms (left) showing Stat3 Y705p and Stat5 Y694p at day 2 for each condition. Representative flow cytometric analysis showing Thy1.1 versus *Foxp3*GFP^+^ expression on day 8 for selected conditions (middle). Quantification (right) showing number of Thy1.1^+^*Foxp3*GFP^+^ at day 8 (n=3, mean ± SEM; ns (nonsignificant), **P<0.01, ****P<0.0001). Statistical differences were tested using one-way ANOVA.

IL-6, a strong inducer of Stat3 activation, can induce low levels of Rorγt expression in splenic Treg cells stimulated *ex vivo*^46^ and has been shown to support Rorγt cells *in vivo*^11^. To evaluate effects of IL-6 on the induciton of effector iTregs *in vitro*, we polarized Treg cells with the graded concentrations of IL-2 and IL-6 (**Fig. 4b**). High levels of IL-2 led to enhanced pStat3, pStat5 and IL-10. In contrast, and in agreement with prior studies, high concentrations of IL-6 led to elevated pStat3 but reduced pStat5 and ablated Treg differentiation. Intriguingly, the frequency of Thy1.1^+^Foxp3^+^ cells was optimized when very low concentrations of IL-6 (0.01ng/mL) were provided in the setting of high-dose IL-2 (50 U/ml; **Fig. 4b**). Polarization in the presence of IL-2 or IL-6 blocking antibodies confirmed that while IL-6 can augment IL-10 secretion by eTreg cells, IL-2 is absolutely required whereas IL-6 opposes the Treg developmental program (**Fig. 4c**). Indeed, even residual IL-6 in serum was sufficient to reduce the total number of Foxp3^+^ cells (**Fig. S4b**). Because a feed-forward mechanism has been described for IL-10R signaling on Treg cells^47^, we also performed IL-10R blocking experiments but found no changes in day 2 pStat3 gMFI or day 8 IL-10 competency (**Fig. S4c**). Blockade of IL-10R *in vivo* also confirmed that IL-10R signaling was not required for IL-10 competent Treg cells in the intestine (**Fig. S4d**). Taken together, these data indicate that IL-2 is necessary and sufficient for the generation of peripherally induced IL-10 competent eTreg cells and that low doses of Stat3 activating cytokines can augment this process in the context of potent Stat5 activation.

### IL-2–mediated Stat3 activation is critical for effector iTreg cell development *in vivo*

Due to pleiotropic effects on both pro- and anti-inflammatory immune cells, IL-2 has been classified as an immune–sensing cytokine that is immune–stimulatory at high doses and immune–regulatory at low doses^48–50^. This differential response of immune cells to IL-2 signaling is attributed to differences in cell type responsiveness and sensitivity^51^; as such, significant efforts have been made to exploit these differences to restrict IL-2 signaling to Treg cells *in vivo* for therapeutic application. IL-2 mutant proteins (muteins) engineered for reduced affinity for γ_c_ preferentially signal on Treg cells *in vivo* due to a lower signaling threshold that reflects their constitutive expression of CD25 (IL-2Ra) compared to other immune cells^52^. IL-2-REH (L18<u>R</u>, Q22<u>E</u>, Q126<u>H</u>) (REH) (**Fig. 5a**) is one such mutein with partial agonist activity that specifically expands existing Treg cells *in vivo* without affecting CD8^+^ T or NK cells^52^. Interestingly, however, this mutein does not induce de novo formation of iTreg cells *in vivo* and, on acute re-stimulation *ex vivo*, induces significantly less *Il10* than WT IL-2^52^. These findings led us to hypothesize that, in contrast to WT IL-2, REH may fail to activate Stat3.

**Fig. 5.**
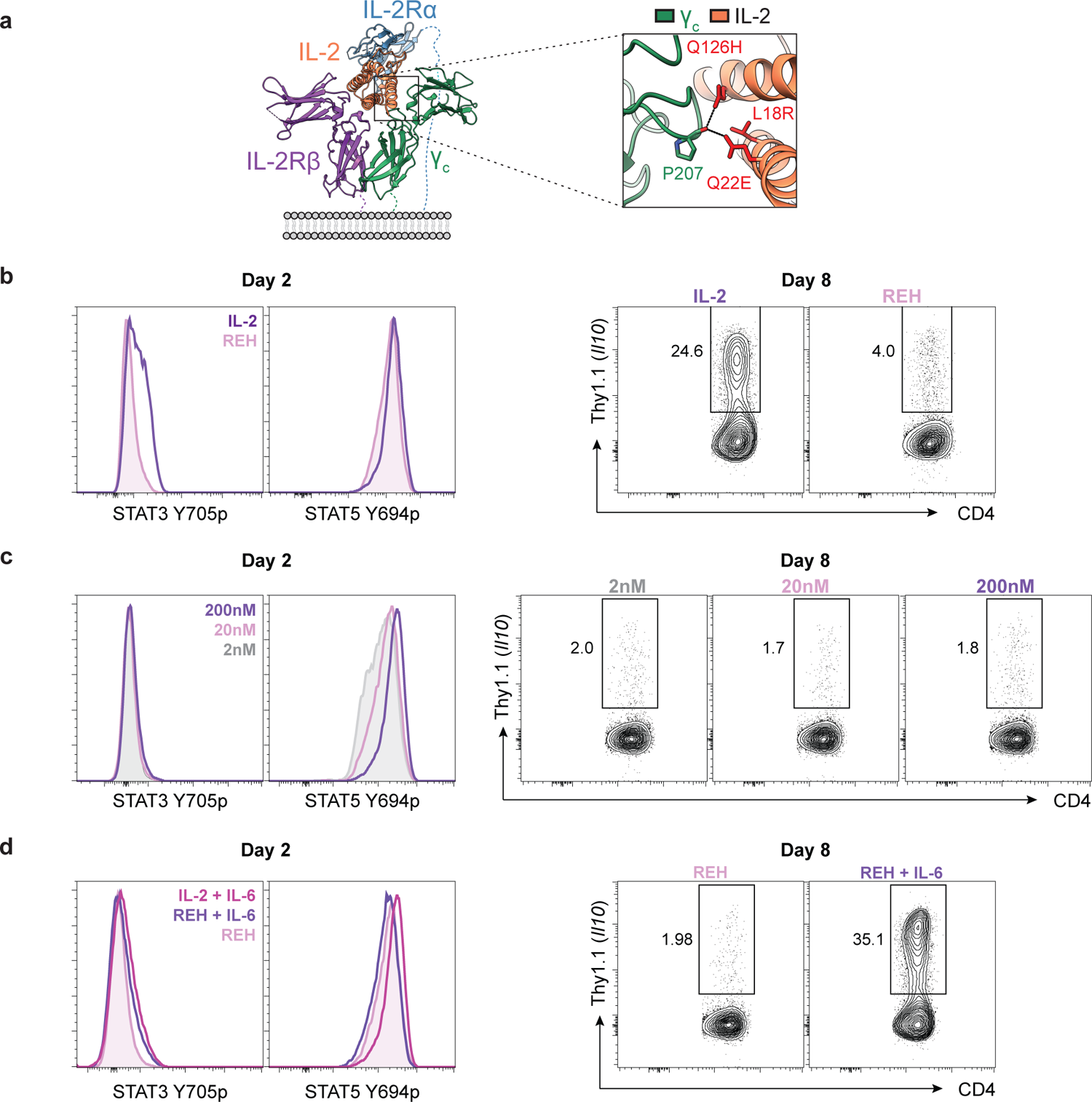
The IL-2 mutein, REH, does not activate Stat3 and fails to induce eTreg cell development. **a**, Schematic showing altered residues of REH that modulate interaction with the IL-2Rγc (green). **b-d**, Naïve CD4 T cells were isolated from 10BiT.*Foxp3*GFP mice and polarized in vitro in iTreg conditions with the indicated cytokines during the induction phase and 50 U/mL IL-2 during re-stimulation. **b,** 200nM MSA-IL-2 (purple) or 200nM MSA-REH (pink). Histograms show Stat3 Y705p and Stat5 Y694p at day 2 for each condition (left). Flow cytometric profiles show Thy1.1 versus CD4 expression (gated on *Foxp3*GFP^+^) at day 8 (right). Experiment performed 3 times. **c,** 2nM (grey), 20nM (pink) or 200nM (purple) MSA-REH. Histograms show Stat3 Y705p and Stat5 Y694p at day 2 for each condition (left). Flow cytometric profiles show Thy1.1 versus CD4 expression (gated on *Foxp3*GFP^+^ at day 8 (right). Experiment performed twice. **d**, 200nM MSA-IL-2 + 0.1ng/mL IL-6 (magenta), 200nM MSA-REH + 0.1ng/mL IL-6 (purple), or 200nM MSA-REH alone (pink). Histograms show Stat3 Y705p and Stat5 Y694p at day 2 for each condition (left). Flow cytometric profiles show Thy1.1 versus CD4 expression (gated on *Foxp3*GFP^+^) at day 8 (right). Experiment performed 3 times. All flow cytometric profiles depict cell number controlled concatenated averages of technical repeats.

To test the actions of REH on Stat3, iTreg cells were polarized with the same concentration of mouse serum albumin (MSA)-cojugates of either REH or WT IL-2 (MSA-REH and MSA-IL-2, respectively; **Fig. 5b**) and pStat3 and pStat5 induction were assessed. It was found that, while pStat5 levels were comparable in cells exposed to either REH or IL-2, pStat3 was detectable in cells activated with MSA-IL-2 but not MSA-REH (**Fig. 5b** and **Fig. S5**). Accordingly, despite restimulation with high-dose WT IL-2 in secondary cultures, IL-10 induction in cells initially activated with MSA-REH showed profund ablation of IL-10 (**Fig. 5b** and **Fig. S5**). The reduction in IL-10 could not be overcome by elevated concentrations of REH, which completely failed to activate Stat3 despite increasing Stat5 phosphorylation (**Fig. 5c**). In agreement with our prior observation that Stat3 activation was dispensible during restimulation, REH was fully capable of inducing Thy1.1 expression in cells polarized in the presence of WT IL-2 (**Fig. S5**). To further validate that the lack of *Il10* induction by REH was dependent upon its inability to activate Stat3, we performed pStat3 rescue experiments. Here, cells were polarized with MSA-REH and low-dose IL-6, which was able to fully rescue Stat3 activation at day 2 (**Fig. 5d**), leading to complete rescue of IL-10 competency at day 8 (**Fig. 5d**). Thus, REH failed to fully support eTreg programming despite comparable Stat5 and iTreg induction and was rescued by Stat3 activation through an independent cytokine receptor (IL-6R), establishing that impaired Stat3 activation by this mutein compromised IL-10 competency. These data reinforce the critical importance of Stat3 activation downstream of IL-2 signaling for the generation of effector iTreg cells *in vitro* and establish that REH has impaired induction of IL-10 due to impaired Stat3 signaling.

To extend these observations *in vivo*, *Stat3^Foxp3^*^Cre^ and *Stat3*^Δ*Foxp3*Cre^ mice were treated with either PBS or MSA-REH (**Fig. 6a**). We found a significant expansion of Helios^+^ but not Helios^-^ Treg cells following treatment with REH; accordingly, there was also a reduction in the ratio of Thy1.1^+^ to Thy1.1^-^ Treg cells, indicating a selective expansion of the non-effector Treg cells (**Fig. 6a** and **Fig. S6a**). There was no significant change in Thy1.1gMFI between groups (**Fig. S6a**). Interstingly, increased total Treg numbers (both effectors and non-effectors) indicated that enhanced proliferation induced by IL-2 treatment is independent of Stat3 activation (**Fig. 6a, Fig. S6a**). Taken together, these data suggest that Stat3 activation downstream of IL-2 dictates colonic effector Treg cell IL-10 competency.

**Fig. 6.**
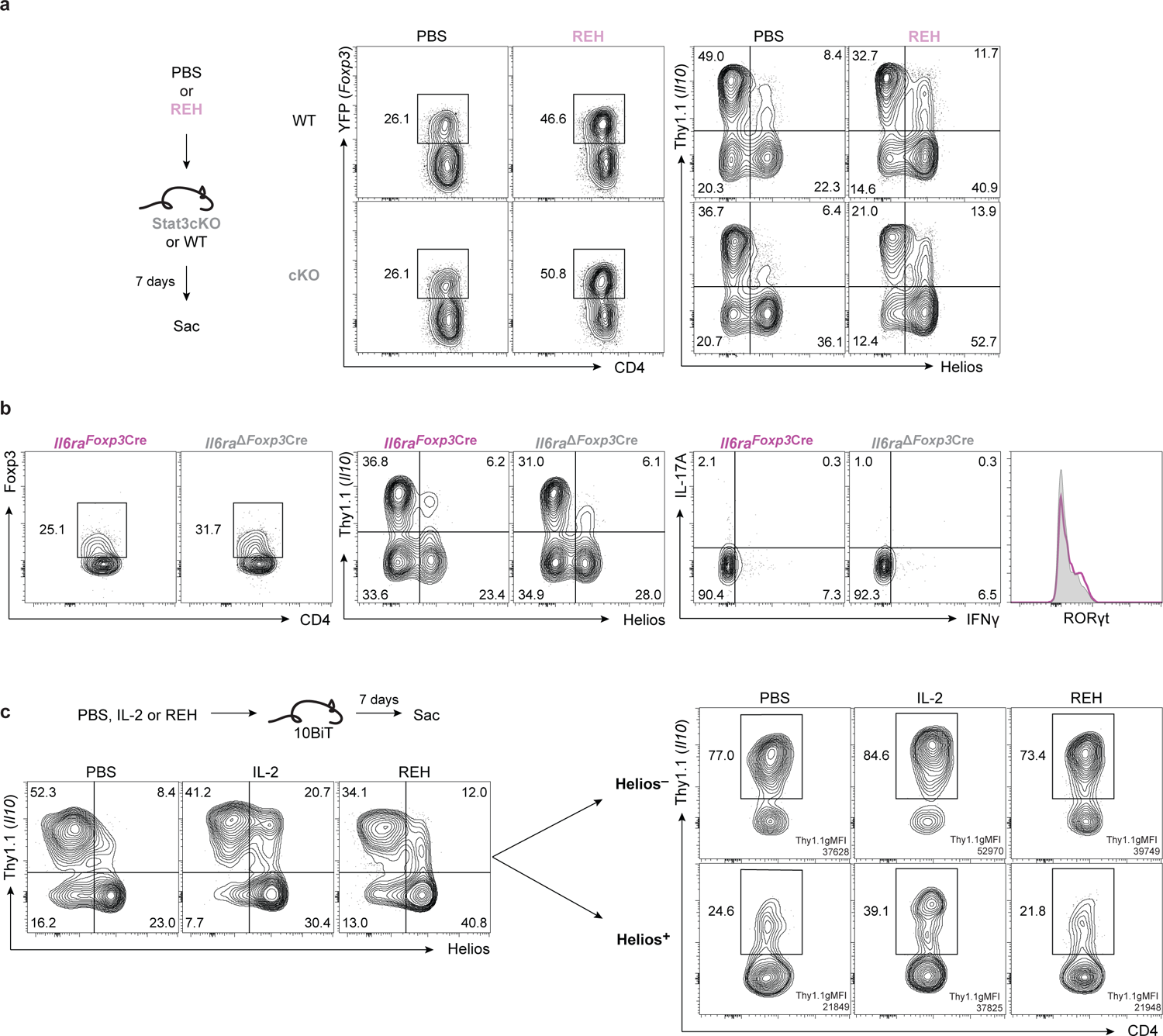
Activation of Stat3 by IL-2 is required for expansion of colonic eTreg cells. **a**, Gender matched *Stat3*^Foxp3Cre^ (WT, black) and *Stat3*^ΔFoxp3Cre^ (cKO, grey) littermate mice were treated I.P. with 30µg/mL of REH (pink) or PBS (black) on days 0, 3, and 6. Animals were sacrificed on day 7 and cells were isolated from the colonic lamina propria. Flow cytometric profiles show *Foxp3*YFP versus CD4 expression (left, gated on CD4^+^TCRβ^+^) and Thy1.1 versus Helios expression (right, gated on CD4^+^TCRβ^+^*Foxp3*YFP^+^). Experiment performed twice. **b**, Colonic lamina propria lymphocytes were isolated from gender matched *Il6raFoxp3*Cre^-^ (WT, purple) and *Il6raFoxp3*Cre^+^ (cKO, grey) littermate mice. Flow cytometric profiles show expression of Foxp3 versus CD4 (left, gated on CD4^+^TCRβ^+^), Thy1.1 versus Helios (middle, gated on CD4^+^TCRβ^+^*Foxp3*YFP^+^), and Il-17A vs Ifnγ (right, gated on CD4^+^TCRβ^+^). Histogram (far right) shows Rorγt expression in Treg cells (gated on CD4^+^TCRβ^+^Foxp3^+^). Experiment performed twice. **c,** 10BiT mice were treated with PBS, 5µg of MSA-IL-2, or 5µg MSA-REH on days 0, 2, 4, and 6 and sacrificed on day 7. Lymphocytes were isolated from the colonic lamina propria and assessed by flow cytometry. Flow cytometric profiles show Thy1.1 versus Helios (left, gated on CD4^+^TCRβ^+^*Foxp3*YFP^+^) and Thy1.1 versus CD4 (top right gated on CD4^+^TCRβ^+^*Foxp3*YFP^+^Helios^-^, bottom right gated on CD4^+^TCRβ^+^*Foxp3*YFP^+^Helios^-^). Thy1.1^+^ gMFI is depicted in for each flow cytometric profile. Experiment performed twice. All flow cytometric profiles depict cell number concatenated averages of biological replicates.

While the foregoing studies indicated that IL-2–induced Stat3 is necessary and sufficient for programming of eTreg cells *in vitro* and *in vivo*, they did not exclude a potentially significant contribution by IL-6. To explore the contribution of IL-6 *in vivo*, *Il6ra* was deleted from Foxp3^+^ T cells by crossing mice carrying a *loxP*-flanked *Il6ra* (*Il6ra^fl/fl^*) allele^53^ with *Foxp3*Cre.10BiT reporter mice (“*Il6ra*^Δ*Foxp3*Cre^”). We found only modest reductions in the frequency of Helios^-^Thy1.1^+^ Treg cells in the *Il6ra*^Δ*Foxp3*Cre^ mice compared to littermate controls and no significant change in the frequency of Helios^+^ Tregs (**Fig. 6b**). Moreover, there was no significant decrement in the ferquency of Rorgt^+^ cell within the Foxp3^+^ population. Accordingly, these mice showed no evidence of spontaneous intestinal inflammation. Similarly, mice with deficiency of *Il6ra* targeted to all T cells (*Il6ra*^Δ*CD4*Cre^) prior to the expression of Foxp3 showed only a similar, modest decrease in Helios^+^Thy1.1^+^ subset and no change in the total number of colonic Foxp3^+^ Treg cells (**Fig. S6b**). Thus, although a contribution for IL-6-induced Stat3 signaling to the development of Helios^+^Thy1.1^+^ iTreg cells could not be excluded, at least at homeostasis its contributions to the development of these cells was minimal, and appeared expendable, supporting a prinicpal role for IL-2 indcued Stat3 signaling for the programming of this population.

To further probe the role of IL-2-mediated Stat3 activation *in vivo*, we treated 10BiT mice with either MSA-REH or MSA-IL-2 (**Fig. 6c**). While both IL-2 variants expanded Helios^+^ Treg cells, REH preferentially expanded Thy1.1^-^ cells while IL-2 expanded Thy1.1^+^ cells (**Fig. 6c**). When we examined the proportion of Helios^-^ and Helios^+^ Treg cells positive for Thy1.1, we found that MSA-IL-2 treatment enhanced not only the frequency of Thy1.1^+^ cells in both subsets but also Thy1.1 expression on a per–cell basis (**Fig. 6c**). MSA-IL-2 treatment also promoted expression of Rorγt in Helios^-^Thy1.1^+^ cells (**Fig. S6c**). Conversely, REH did not affect the frequency or gMFI of Thy1.1, or Rorγt expression, in either subset (**Fig. 6c, Fig. S6c**). Taken together, these data indicated that, at homeostasis, IL-2–mediated Stat3 activation is necessary and sufficient for development of IL-10– competent Rorγt iTreg cells in the colon.

### IL-2 and pStat3 modulate cis-regulatory elements within *Il10* early in iTreg development

To examine mechanism(s) underlying IL-2-mediated Stat3-dependent induction of *Il10*, we examined differential chromatin accessibility (DCA) at day 2 in CD4 T cells polarized with or without IL-2 and in WT versus Stat3-deficient T cells using transposase-accessible chromatin with high-throughput sequencing (ATAC-seq, **Fig. 7a, b**). Multiple genomic regions that correlated with conserved non-coding sequences (CNSs; mouse-human) gained accessibility across the *Il10* gene locus contingent on IL-2 signaling (**Fig. 7a**) Notably however, no major differences in accessibility were evident in Stat3-deficient cells (**Fig. 7b**). Genome–wide motif analysis of DCA peaks showed that a large proportion of motifs enriched in the Stat3 cKO v WT ATAC-seq were also enriched in the IL-2 ATAC-seq (**Fig. 7c**). The shared motifs included Stat5, Stat3, and TFs previosly reported to be important in the regulation of *Il10*, including PRDM1^34^ and bHLHE40^54,55^ (**Fig. 7d**). These data indicated that while IL-2 signaling exerted pronounced early effects on chromatin accessibility across the *Il10* locus, this was largely indpendent of Stat3.

**Fig. 7.**
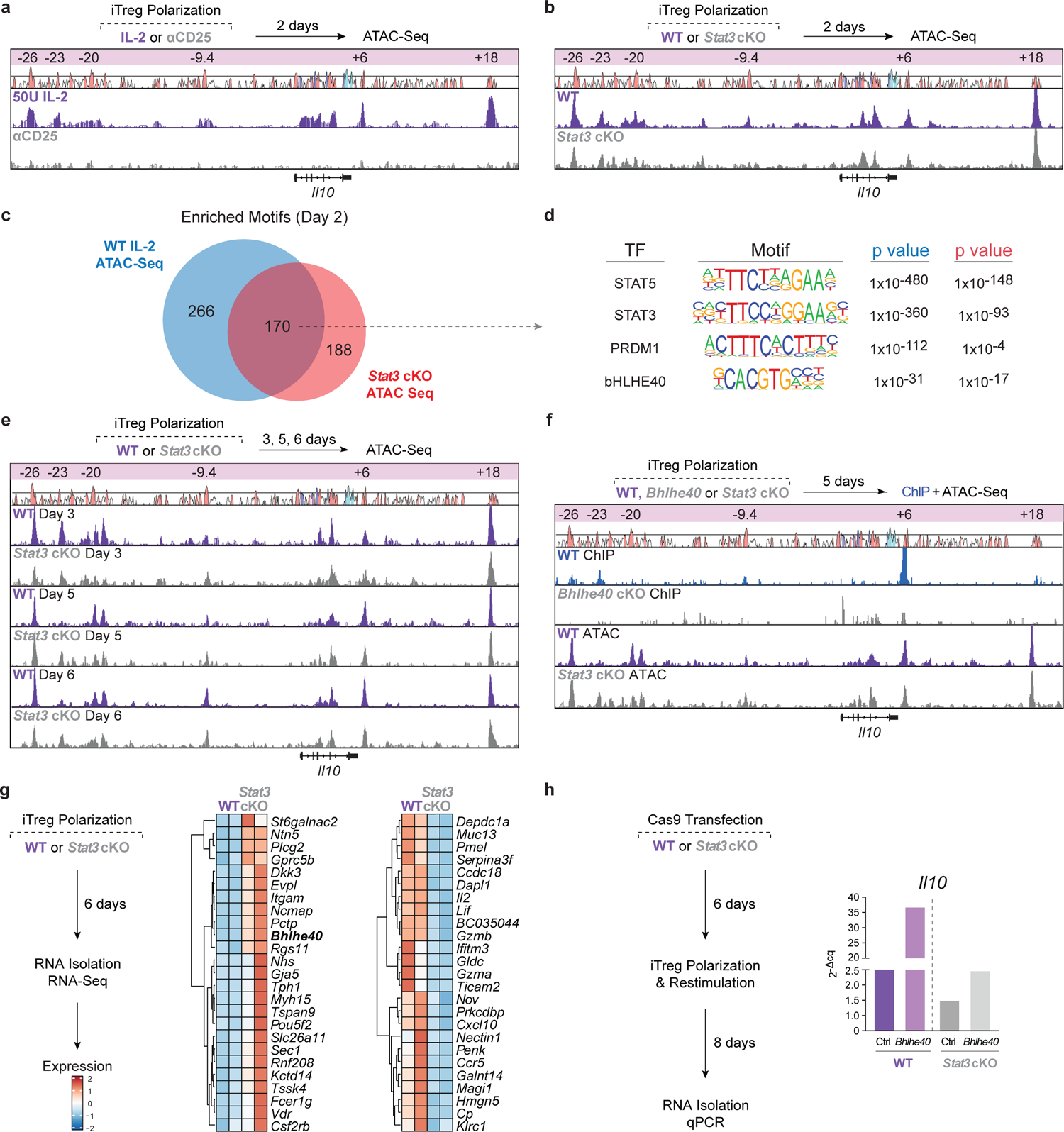
Stat3 and IL-2 dictate chromatin accessibility at *Il10* early in iTreg cell development. **a**, Naïve CD4^+^ T cells were polarized as in Fig. 3a with 50U/mL IL-2 (purple) or 0U/mL IL-2 + 10 µg/mL αCD25 (grey). ATAC-seq was performed on live cells on day 2. Representative ATAC-Seq tracks show differential accessibility (DA) within the *Il10* gene body at intron 4-SRE and downstream at +18kb (magenta asterisks). **b,** Naïve CD4^+^ T cells were isolated from *Stat3*^CD4Cre^ (WT, purple) or *Stat3*^ΔCD4Cre^ (cKO, grey) and polarized as in Fig. 3a with 50U/mL IL-2. ATAC-seq was performed on live cells on day 2. Representative ATAC-Seq tracks show no significant DA across the *Il10* locus. **c,** Venn diagram displaying the overlap between genome-wide motif enrichment analysis of differentially accessible regions in the ATAC-Seq data from **a** (blue) and **b** (red). **d,** Table showing a selection of shared enriched motifs, their associated transcription factors, and p value for each experiment. **e,** Naïve CD4^+^ T cells were isolated from *Stat3*^CD4Cre^ (WT, purple) or *Stat3*^ΔCD4Cre^ (cKO, grey) and polarized as in Fig. 3a with 50U/mL IL-2. ATAC-seq was then performed on live cells on days 3, 5, and 6. Naïve CD4^+^ T cells were also isolated from C57BL/6 WT mice (black) and polarized as in Fig. 3a with 50U/mL IL-2. pStat3 CUT&RUN was then performed on days 3, 5, and 6. Magenta asterisks shown below the tracks denote DA between Stat3 WT and cKO iTreg cells (ATAC-Seq) or pStat3 binding regions (CnR). Data represent the combined analysis of two biologically independent samples.

To further parse out the effect of Stat3 on the chromatin landscape downstream of IL-2 signaling, we performed kinetic ATAC-Seq in iTreg cells polarized *in vitro* from *Stat3*^Δ*CD4*Cre^ and *Stat3^CD4^*^Cre^ naïve CD4 T cells. During the polarization phase (d2, d3, d5), Stat3 exerted only modest effects on accessibility at CNSs across the *Il10* locus, primarily limited to sites –23kb (CNS-23; d5) and +18kb (CNS+18; d3) upstream and downstream of the gene start site, respectively (**Fig. 7e**). Though Stat3-dependent remodeling of the *Il10* locus was limited, accessibility of multiple regions was modulated throughout iTreg development and maturation. CNS-23 and CNS+18 were opened early (d2) following IL-2 signaling but became less accessible over the time course (**Fig. 7b,e**). CNS-26, CNS-20, and introns 3 and 4 were similarly remodeled by d3 but maintained a relatively constant degree of accessibility while CNS+6 gained in accessibility as iTregs acquired IL-10-competency. Taken together, these data indicated that multiple conserved non-coding elements in the *Il10* locus are differentially remodeled during iTreg development to promote *Il10* expression and while IL-2–pStat3 was important in establishing *Il10* transcriptional competency early in iTreg development, direct actions on the locus following restimulation were limited. These data are congruent with our *in vitro* assays, where we found that early exposure to IL-2–pStat3 dictated *Il10* transcriptional competency (**Fig. 4**), whereas cytokine exposure on re-stimulation had minimal additional effect (**Fig. S5a).**

Because Stat5 is the dominant output of IL-2 signaling, we performed CUT&RUN-seq in *Stat3*^Δ*CD4*Cre^ and *Stat3^CD4^*^Cre^ *in vitro* iTreg cells to evaluate Stat5 binding in the presence and absence of Stat3. Bioinformatic analyses found enriched binding of Stat5 to the gene body of *Il10* at day 2, the promoter at day 5, and CNS+18 on days 3, 5, and 6 in WT cells (**Fig. S7b**). Further, we found that the absence of Stat3 had minimal effect on Stat5’s ability to bind the locus (**Fig. S7b**), indicating that Stat3 is unlikely to promote *Il10* transcriptional competency through direct effects on Stat5 binding, whether positive or negative. Thus, although these data due not preclude cooperative actions of Stat5 and Stat3 at the *Il10* locus, Stat5 is not dependent on Stat3 for binding nor is it clearly antagonized by Stat3 at *Il10*.

### Stat3 regulates the transcriptional repressor bHLHe40

Motif analyses of the regions identified by ATAC-seq and pStat5 CUT&RUN-seq identified predicted binding sites for a number of TFs (**Fig. S7a-b**). One of these, basic helix-loop-helix e40 (bHLHe40) – a negative regulator of *Il10*, was predicted to bind at CNS-23, a site that lost accessibility during iTreg development and was negatively regulated by Stat3, as indicated by increased DCA in the absence of Stat3 compared to WT (**Fig. 7e**). CNS+18, which similarly had diminished accessibility late also contained a bHLHe40 consensus binding motif. bHLHE40 has been reported to inhibit *Il10* in CD4 T cells, including Th1 and Tr1 cells, by binding to various sites at *Il10*^54,55^. Because Stat3 negatively regulated accessibility of a putative Bhlhe40–binding region at *Il10*, we wanted to understand whether this mechanism was important in Stat3-mediated regulation of IL-10 competency.

To validate bHLHe40 binding to *Il10* CNS-23, we analyzed previously published Bhlhe40ChIP-seq data in Th1 cells^54^ and confirmed recruitment of bHLHe40 to CNS+6 but also found substantial peak enrichment at CNS-23 and to a lesser degree, at CNS-26, −9.4 and CNS+18 (**Fig. 7f**). To understand whether Stat3 regulated the expression of *Bhlhe40*, RNA-sequencing of *in vitro* derived iTreg cells from *Stat3*^Δ*CD4*Cre^ and *Stat3^CD4^*^Cre^ mice was performed. We found that, in absence of Stat3 signaling, *Bhlhe40* transcript was significantly upregulated, indicating that Stat3 may be a negative regulator of *Bhlhe40* (**Fig. 7g**). These data indicated a potential two-pronged mechanism of Stat3’s regulation of bHLHE40 – (1) reduced accessibility of bHLHe40 binding sites in the *Il10* locus and (2) reduced expression of *Bhlhe40* itself. To understand whether this mechanism played a major role in Treg cell IL-10 competency, CRISPR-mediated knockdown of *Bhlhe40* was performed in iTregs deficient for Stat3 differentiated *in vitro* (**Fig. 7h**). We found that when *Bhlhe40* expression was reduced in WT or Stat3 deficient cells, IL-10 competency was enhanced (**Fig. 7h**). However, the increase in *Il10* expression in WT cells was significantly greater than in Stat3-deficient cells, implicating Stat3’s regulation of bHLHe40 as one, albeit not the only, mechanism underlying Stat3– mediated *Il10* expression in iTreg cells.

## Discussion

Here, we have elucidated a new role for IL-2 in Treg cell biology, adding to the known functions of IL-2 a central role for programming Helios^-^ iTreg cells for eTreg cell-associated IL-10 competency. Remarkably, IL-2-induced Stat3, not Stat5, activation was key to this programming, acting early in iTreg cell development to promote IL-10 expression following further antigen-driven differentiation. These findings expose a heretofore unappreciated role for the dual Stat5-Stat3 output of the IL-2 receptor with implications for mechanisms that control *Il10* transcription, and provide a new perspective on the interplay of these two Stats in regulating the iTreg-Th17 developmental axes. Our findings also expose a more nuanced view on the complementary and competing roles of these T cell subsets in maintaining immune homeostasis, particularly in the gut where these populations dominate. Because IL-10-producing eTregs are a major cell population contributing to maintenance of homeostasis, appropriate ratioing of Stat5/Stat3 assumes critical new importance in the biology of IL-2-mediated immune regulation.

The intestines and its associated lymphoid tissues house large numbers Treg cells with distinct ontologies and functions; several subsets of tTreg and iTreg cells that are developmentally and fuctionally distinct coexist in the large and small intestines^58^. Results herein demonstate a clear distinction between Helios^+^ and Helio^-^ Treg cells in the large intestine (LI), whether at homeostasis or pathobiont colonization: Treg cells with active expression of *Il10* are primarily limited to the Helios^-^ iTreg compartment, which are predominantly Rorγt^+^ and essential to maintenance of gut homeostasis, whereas the Helios^+^ tTreg cells are devoid of active *Il10* expression^58^. While this could indicate fundamental differences in the programming of the major Treg subsets that reflect thymic versus post-thymic origins, the delivery of exogenous WT IL-2, but not the IL-2 mutein REH, induced a substantial fraction of Thy1.1^bright^ Helios^+^ Treg cells in the colon, indicating the capacity of this subset to acquire expression of IL-10 driven by IL-2 when Stat3 output of the IL-2R was engaged. Although our studies did not discriminate between intrathymic versus extrathymic effects of exogenous IL-2 administration, if the actions of IL-2-induced Stat3 in licensing iTreg cells for IL-10 competency have direct parallels with tTreg cell programming, i.e., during inital induction of Foxp3, exogenous IL-2 may have acted principally on tTregs developing in the thymus that would otherwise not have been adequately “primed” for subsequent eTreg cell development. If so, given the low level of IL-10 production among tTreg cells in the LI, this implies a difference in IL-2 signaling intensity and/or Stat3 coupling during the development of tTreg versus iTreg cells. The role of IL-2-induced Stat3 in driving IL-10 production by tTreg cells will require further study.

Implementation of a robust, two-stage system for the development of stable, IL-10^+^ eTreg cells *in vitro* allowed us to dissect the timing of signals required for their programming. The window for induction of IL-10 competency was surprisingly narrow and limited to the early phase of development, coinciding with initial IL-2-driven signaling required for iTreg cell differentiation. The maximal effect of Stat3 signaling inhibition was exerted over the initial 3 days of differentiation and progressively attenuated thereafter; it was nearly completely lost if deferred to the restimulation phase. Although additional studies will be needed to define exactly how rigid this early Stat3 signaling window is for aquisition of IL-10 competency, i.e., the degree to which Treg cells can acquire IL-10 expression after failing to receive Stat3 imprinting during initial induction, findings herein would appear to place substantial constraints on the trajectory of full eTreg cell development that are deterministic early in iTreg commitment.

This system also revealed an absence of IL-10 expression by iTreg cells during the inductive phase; iTregs failed to express IL-10 unless reactivated for at least 2-3 days in secondary cultures supplemented with exogenous IL-2. Thus, unlike pro-inflammatory effector CD4 T cells (e.g., Th17, Th1), which are competent for effector cytokine expression either late during the inductive phase or immediately upon reactivation with antigen, our findings indicate that iTreg cells require a period of additional maturation following reactivation to express IL-10. Deferred acquisition of IL-10 production by developing eTreg cells and a reliance on IL-2 provided by Teff cells in non-lymphoid tissues for this maturation ensures that IL-10 does not cripple host-protective Teff cell responses prematurely, and couples the terminal differentiation of eTregs to the inflammatory micrenvironment. In essence, this provides a second window to sense and further differentiate – or not – to acquire greater repressive functionalities contingent on, and under the influence of, IL-2 sourced from pro-inflammatory Teff cells. However, in contrast to the inductive phase, during which Stat3 output of the IL-2 receptor was critical, IL-2 signaling required for completion of eTreg programming during this second, maturational phase appears to be Stat3-independent. The uncoupling of Stat5 and Stat3 signaling provided by REH should enable additional in-vivo studies to further explore this issue.

Importantly, IL-2 was both necessary and sufficient for robust IL-10 expression by eTreg cells. Although augmented pStat3 signaling activated through an independent cytokine receptor (IL-6) enhanced IL-10 induced by saturating IL-2 *in vitro*, the effect was relatively modest. Moreover, T cell- or Treg-targeted deficiency of IL-6 signaling had minimal impact on Treg cell expression of IL-10 in the LI and, conversely, IL-2 supplementation increased their numbers. Thus, IL-2 signaling alone appears sufficient to underpin the homeostatic network that buffers microbiota-driven inflammation in the large intestine. Accordingly, while other Stat3-activating cytokines, such as IL-6, may contribute to the programming of IL-10-competent eTreg cells – perhaps under conditions of limited IL-2 availability – at homeostasis this would appear to be dispensable. Signaling outputs of non-IL-2 cytokine receptors distinct from Stat3 also appear to be dispensable.

Notably, however, in a companion study (Dean *et al*. submitted), we have identified a mechanism by which co-signaling through the TNF receptor superfamily (TNFRSF) member DR3 (TNFRSF25) potentiates the programming of effector iTreg cells induced by limiting quantities of IL-2. Shown previously to upregulate components of the IL-2R complex to enhance Treg cell numbers^59,60^, we now find that DR3 signaling also promotes the increased amplitude and duration of Stat3 signaling activated by suboptimal IL-2 in developing iTreg cells, thereby enhancing IL-10 competency. Expressed rapidly following antigenic activation of naive CD4 T cells^61,62^, DR3 is available to supplement IL-2-induced Stat3 signaling contingent on availability of its ligand, TL1A (TNFSF15), which is primarily expressed by inflammatory myeloid cells^63,64^. Because DR3 signaling does not directly activate Stat3, this implicates an indirect mechansim by which this pathway augments Stat3 signaling initiated by IL-2. A previous report identified a role for induction of the NF-κB family member RelA by TNFRSF members in supporting the survival of eTreg cells^65^, which deserves further study. Interestingly, DR3 is unusual among TNFRSF members in also being constititively expressed on “resting” iTreg cells^65^, making it uniquely positioned to amplify iTreg and eTreg cell numbers under inflammatory conditions, thereby providing a layer of regulatory control by which the IL-2–sensing regulatory network may be “tuned” by auxilliary cytokine signals.

To begin to explore mechansisms by which Stat3 contributes to eTreg differentiation, we focused on the *Il10* locus. While IL-2 signaling had a major impact on chromatin accessibility at the *Il10* locus, this appeared to be largely due to Stat5-dependent actions; effects of Stat3 signaling were more nuanced. Although our results do not preclude a role for Stat3 in directly modulating chromatin accessibility at the *Il10* locus, its non-redundant role in regulating the *Il10* locus would appear to be contingent on the induction of additional *trans* factors that are likely to cooperate with Stat3 to control *Il10* transcription. One candidate factor is Rorγt, expression of which is strictly dependent on Stat3 and which has a strong correlation with IL-10 competency in eTreg cells in the gut, yet we nevertheless find a substantial fraction of Helios^-^IL-10^+^ iTreg cells lack Rorgt expression. While additional studies will be needed to fully map early actions of Stat3 signaling that contribute to *Il10* expression competency, at least one mechanism involves Stat3-dependent repression of the transcriptional repressor, Bhlhe40, acting through the cis-regulatory element, CNS-23, although clearly additional mechanisms are at play. In view of the technical challenges posed by limiting cell numbers for mapping transfactor networks by pan-genome binding, implementation of the in vitro system reported herein for the generation of iTreg and eTreg cells in adequate numbers and under well-defined conditions should prove valuable.

Our findings have significant implications for IL-2–based therapeutics. While tremendous strides have been made to harness the potential of engineered cytokines for immune-mediated disease indications^31^, clinical efficacy remains a barrier^66^. The discovery of an unappreciated role for IL-2 in driving Stat3 phosphorylation in developing iTreg cells, and the inability of at least one synthetic IL-2 variant to activate Stat3, suggests an incomplete understanding of IL-2 biology remains a liability for some clinical applications and highlights the potential importance of preclinical studies that encompass a broader assessment of immune modulatory effects of IL-2 variants. Insofar as IL-10– producing eTregs may represent highly effective agents for reversal of immune-mediated diseases, particularly in inflammatory bowel diseases – and especially in the clinical settings of ongoing T cell-driven inflammation – appropriate consideration of the balance of Stat5-Stat3 outputs would appear to be warranted. This extends to use of these agents for optimal ex-vivo generation of Treg cells for adoptive cell-based therapies^67^. Conversely, while IL-2 muteins with attenuated Stat3 activities could prove problematic for therapies that target immune-mediated disease, they could benefit IL-2–based efforts to augment CD8 responses in tumor microenvironments where Treg cells may contribute to T cell exhaustion that limits anti-tumor cytotoxic activity.

## Acknowledgments

The authors thank Drs. Hui Hu, Charles Elson, Craig Maynard (University of Alabama Birmingham (UAB), Robin Lorenz, (Genentech), and members of the Weaver Laboratory for their helpful comments and suggestions. We thank La Jolla Institute for sequencing assistance with gene expression studies. We gratefully acknowledge Brenda Dale, Emma Higginbotham, and Kim Nguyen for their technical expertise and Elaina Harris for editorial assistance. We also thank the UAB Comprehensive Flow Cytometry Core for flow cytometric sorting and single cell RNA-sequencing library preparation assistance. This work was supported by an NIH R01s DK115172 and AI161717 (C.T.W. and R.D.H), an NIH training grant F30 5F30DK121398-04 (E.C.D), and the UAB Medical Scientist Training Program T32GM008361 (E.C.D.).

## Methods

### Animals

All animals were on a C57BL/6 background. B6.129S1-Stat3tm1Xyfu/J (*Stat3* floxed), B6.129(Cg)-Foxp3tm4(YFP/icre)Ayr/J (*Foxp3*YFPCre), B6.Cg-Tg(Cd4-cre)1Cwi/BfluJ (*Cd4*Cre), C57BL/6, B6.129S7-Rag1tm1Mom/J (*Rag1*^-/-^), and C57BL/6-Il17atm1Bcgen/J (*Il17a*eGFP) were purchased from The Jackson Laboratory (strain numbers: 016923, 016959, 022071, 002216, and 018472). 10BiT (Tg(Il10-Thy1^a^)1Weav)^16^ were generated and bred at the University of Alabama at Birmingham (UAB) animal facility. *Foxp3*GFP (fusion) mice were a gift from Alexander Rudensky (Memorial Sloan Kettering Cancer Center), B6.Cg-Foxp3tm1Kuch/J (*Foxp3-*GFP (IRES)) mice were a gift from Vijay Kuchroo (Brigham and Women’s Hospital), B6;SJL-Il6ratm1.1Drew/J (*Il6ra* floxed) mice were a gift from Angela Drew (University of Cincinnati), *Rorc*eGFP mice were a gift from Gerard Eberl (Pasteur Institute), and *Il2*Cre mice a gift from Andre Ballasteros-Tatos (UAB). *Stat3* floxed, 10BiT and either *Foxp3*YFPCre or *Cd4*Cre mice were intercrossed to generate *Stat3*fl/fl.*Foxp3*YFPCre.10BiT or *Stat3*fl/fl.*CD4*Cre.10BiT mice. 10BiT and either *Foxp3*GFP or *Foxp3*-GFP mice were intercrossed to generate 10BiT.*Foxp3*GFP or 10BiT.*Foxp3-*GFP mice dual reporter mice. 10BiT and *Rorc*eGFP mice were intercrossed to generate 10BiT.*Rorc*eGFP dual reporter mice. *Il6ra* floxed, *CD4*Cre, and 10BiT mice were intercrossed to generate *Il6ra*fl/fl.*CD4*Cre.10BiT mice. *Il2*Cre and 10BiT mice were intercrossed to generate *Il2*Cre.10BiT mice.

Animals were bred and maintained under specific pathogen-free conditions in accordance with UAB Institutional Animal Care and Use Committee regulations. All studies were performed on gender-matched mice 8-14 weeks of age. For Foxp3 conditional deletion, mice heterozygous for the floxed allele and homozygous/hemizygous for *Foxp3*YFPCre were bred to ensure all offspring carried the *Foxp3*YFP reporter allele. For CD4 conditional deletion, mice homozygous for the *Il6ra* floxed allele and heterozygous for the *CD4*Cre allele were bred with mice homozygous for the *Il6ra* allele but negative for *CD4*Cre.

### Staining antibodies and flow cytometry

Surface staining was performed in PBS with 2% FBS for 15 min at 4°C. Intracellular staining for transcription factors and cytokines was performed using the eBioscience Foxp3 staining kit or the BD Cytofix/Cytoperm kit (eBiosciene 00-5523-00, BD 554714). For phospho-Stat flow cytometry, cells were fixed in 1.6% PFA for 10 min at room temperature, permeabilized in 100% MeOH for 30 min on ice, and then stained for 1 hour in 2% FBS at room temperature. Unless otherwise noted in figure legends, cells were not restimulated before phospho-Stat staining. All flow cytometry data were acquired on an Attune NxT (Thermo Fisher Scientific) and analyzed with FlowJo software (Tree Star, Eugene, Oregon). In experiments staining for IL-10, IL-17A or IFNγ, cells were re-stimulated for 3 hours at 37°C in PMA/ionomycin before staining.

### Cell isolation

Mice were sacrificed using isoflurane prior to removal of spleens and colons. Spleens were manually disrupted in complete RPMI-2 using frosted glass slides. RBCs were lysed using Ack lysis buffer (Fisher 50983219), washed, and the remaining cells were resuspended in RPMI-2 or RPM-10. Colon tissue was flushed with HBSS-2 and cut longitudinally. If colons were used for histological purposes, they were cut length wise in half and then one half was cut into three sections for the proximal, middle, and distal colon and fixed in 10% (wt/vol) formalin. The other half was then used to isolate lymphocytes. Colonic sections utilized in lymphocyte isolation were cut into half centimeter long strips and were then manually disrupted with scissors. The tissue was then incubated with gentle magnetic stirring for 45 minutes at 37°C with collagenase IV (Millipore Sigma C5138-5G) with DNase (1 mg/mL, Sigma) in RPMI-10. Cells were purified by Percoll gradient (40%/75%, 20 min, 600g, 25°C).

### In vitro T cell activation

Naïve CD4^+^ T cells were magnetically enriched from total splenocytes using negative selection (Miltenyi Naïve CD4^+^ T Cell Isolation Kit 130-104-453) on LS Columns (Miltenyi 130-042-401). For standard iTreg polarizations, naïve T cells were stimulated for 5 days in 96-well flat-bottom plates (Corning 3596) coated with 2.5ug/mL anti-CD3 (UAB). Cells were stimulated in complete RPMI-10 supplemented with 50U/mL hIL-2 (Roche 11011456001), 2.5ng/mL TGFβ (R&D 240-B-10), 10nM retinoic acid (Sigma R2625-500MG), 100ug/mL L-ascorbic acid (Sigma A4404-100MG), and 2ug/mL aCD28 (Fisher BDB553295). On day 5, cells were moved to new 96-well flat-bottom plates (Cornig 3596) coated with 2.5ug/mL aCD3 and re-stimulated for an additional 3 days in complete IMDM-10 supplemented with 50U/mL IL-2 (Roche 11011456001). For Th17 polarizations, CD4^+^ T cells were isolated via positive magnetic selection according to the manufacturer’s instructions (Invitrogen 11331D). Cells were polarized with irradiated CD4-depleted feeders (1:10, 320Gy 12.5 mA for 10 minutes) for 6 days in 96-well flat-bottom plates in complete RPMI-10 supplemented with 2.5ug/mL aCD3, 20ng/mL IL6 (R&D 406-ML-005), 2.5ng/mL TGFβ (R&D 240-B-10), 5ug/mL aIFNγ (UAB XMG), and 5ug/mL aIL-4 (UAB 11B11).

RPMI-10 and IMDM-10 contained RMPI (RPMI Fisher MT10-040-CM) or IMDM (IMDM Corning MT10016CV) and were supplemented with bovine calf serum (10%, HyClone), 1mM sodium pyruvate (Corning), 2mM l-glutamine, nonessential amino acids (Corning), 100 IU/mL penicillin, 100ug/mL streptomycin, and 50uM 2-mercaptoethanol (Sigma). RPMI-2 contained RPMI and was supplemented with bovine calf serum (2%) and penicillin-streptomycin. HBSS-2 contained HBSS (Fisher MT-21-021-CV) and was supplemented with bovine calf serum (2%) and penicillin-streptomycin.

### RNA-Seq analysis

For sample preparation and hybridization in Fig. 2, colonic Foxp3(GFP)^+^IL-10(Thy1.1)^+^ and Foxp3(GFP)^+^IL-10(Thy1.1)^-^ cells were sorted into Trizol and total RNA was isolated with miRNeasy Micro Kits (QIAGEN 217084) according to the manufacturer’s instructions. For each experiment, 5 mice were combined to yield sufficient cells for RNA isolation. Libraries were prepared at La Jolla Institute for Immunology using Illumina Stranded mRNA Prep, Ligation kit (20040532) and were sequenced using an Illumina NovaSeq with paired end 50 reads. For sample preparation and hybridization in Fig. 8, *in vitro* derived Foxp3^+^ cells were harvested on day 2 and day 6 into Trizol and total RNA was isolated with miRNeasy Micro Kits (QIAGEN 217084) according to the manufacturer’s instructions. Libraries were prepared at La Jolla Institute for Immunology using SMARTer Stranded RNA-Seq kit (Takara 634962) and were sequenced using an Illumina HiSeq2500 in Rapid Run Mode with paired end 50 reads.

**Fig. 8.** Stat3 regulates the activity of bHLHE40. **a**, Representative bHLHE40 ChIP-Seq and Stat3cKO ATAC-Seq tracks. bHLHE40 Chip-Seq data in Th1 cells analyzed from previously published data from Huynh et. al (citation 54 in this version). For ATAC-Seq data, naïve CD4^+^ T cells were isolated from *Stat3*^CD4Cre^ (WT, purple) or *Stat3*^ΔCD4Cre^ (cKO, grey) and polarized as in Fig. 3a with 50U/mL IL-2. ATAC-seq and was then performed on live cells on days 3, 5, and 6. Magenta asterisk denotes bHLHE40 binding enrichment (ChIP-Seq) and DA between Stat3 WT and cKO iTreg cells (ATAC-Seq) at day 5. **b,** Naïve CD4^+^ T cells were isolated from *Stat3*^CD4Cre^ (WT, purple) or *Stat3*^ΔCD4Cre^ (cKO, grey) and polarized as in Fig. 3a with 50U/mL IL-2. ATAC-seq was then performed on live cells on days 3, 5, and 6 (only day 5 shown here). Naïve CD4^+^ T cells were also isolated from C57BL/6 WT mice (black) and polarized as in Fig. 3a with 50U/mL IL-2. pStat3 CUT&RUN was then performed on days 3, 5, and 6. Magenta asterisks denote DA between Stat3 WT and cKO iTreg cells at *Bhlhe40*, as well as sites of pStat3 binding at *Bhlhe40*. **c,** Naïve CD4^+^ T cells were isolated from *Stat3*^CD4Cre^ (WT) or *Stat3*^ΔCD4Cre^ (cKO) and polarized as in Fig. 3a with 50U/mL IL-2. On day 6, live CD4^+^ cells were subjected to RNA-Sequencing. Heatmap depicts relative expression (z-score) of differentially expressed genes between WT and cKO iTreg cells. **d,** Workflow of Crispr-mediated *Bhlhe40* knockdown (left) – CD4^+^ T cells were isolated from Stat3 WT or cKO spleens and transfected with *Bhlhe40* or control (ctrl) guide RNAs (sgRNAs). 6 days after transfection, cells were polarized in iTreg conditions with 50U/mL of IL-2 as described in Fig. 3a. On day 8, cells were collected and processed for RNA isolation. Bar graph shows percent change in *Il10* expression. Data represent the combined analysis of two biologically independent samples.

### Pre-processing of bulk RNA-sequencing data

Single-end or paired-end reads with lengths of 50 nucleotides (∼20M reads per condition) were generated for subsequent bioinformatics analysis. Adaptors were trimmed and aberrant reads were removed using TrimGalore (v.0.4.5). The quality controlled reads were mapped onto the mouse genome build GRCm38 (ENSEMBL.mus_musculus.release-75) using STAR (v.2.5.3)^68^. BAM files were sorted using SAMtools (v. 0.1.18)^69^, and reads were counted for each gene using HTSeq (v.0.7.2)^70^. Quality of libraries was assessed by overall mapping rate, and libraries with less than 70% mapping rate were discarded from further analysis.

### Exploratory analysis and quality control

The R software environment (version 3.6.1) was used for statistical analysis. Gene counts were filtered for genes catalogued by ENSEMBL (ENSEMBL.mus_musculus.release-75) as protein_coding. We subjected cleaned sequencing data to a battery of exploratory analyses using the R package DESeq2 (v.1.18.1)^71^ in order to inform model selection for analysis of differential expression, including heatmaps, distance matrices, and principal component plots of the top 500 differentially expressed genes after rlog transformation^71^. We visually inspected these together in order to identify low-quality samples, outliers, and unexpected trends relating to batch ID or other important covariates to be used in modeling.

### Differential Gene Expression Analysis

Differential expression analysis was performed using DESeq2 (v.1.18.1)^71^ using R (v.3.6.1). Dispersion shrinkage of fold changes was performed with the ASHR algorithm^72^. Briefly, Wald Tests were selected in place of likelihood ratio tests and stratified analysis outperformed pooled analysis for most models. Final model specifications for each of these tests are available by reasonable request. We set alpha, our allowable type I error rate, at a 0.05 after correction for multiple testing using independent hypothesis weighting^73^. For each model, we assessed model fit and model validity based on dispersion estimates, overall type I error rate, Cooks’ distances, inspection of count-dispersion plots and MA plots (log2 fold change versus log2(base mean) for competing models for each research question. The rlog() function from the DESeq2 package was utilized for variance stabilization and was applied to all samples. These data were z-scored and visualized with heatmaps via ComplexHeatmap (v2.11.1) package in R. The “RdBu” color palette was obtained from the R package RColorBrewer (v.1.1.2) and was expanded to 1,000 shades using the R function colorRampPalette().

### Gene Set Enrichment Analysis

The fgsea R package (v.1.4.0)^74^ was used for gene set enrichment. Gene sets used were MSigDB Hallmark gene sets^75–77^. The input for GSEA was the regularized log expression values obtained from DESeq2 which was filtered to remove genes with mean expression ≤ 0. A p-value quantifying the likelihood that a given gene set displays the observed level of enrichment for DE genes was calculated using Fast Gene Set Enrichment Analysis (fgsea, v.1.4.0) with 1 million permutations. Gene set enrichment p-values of Normalized Enrichment Scores (NES) were corrected with the Benjamini-Hochberg procedure^78^. The top enriched terms were visualized by the lollipop plots or dot plots using R package ggplot2(v3.3.5).

### ATAC-Seq and motif analysis

ATAC-Seq was performed as described^79^. Following sequencing and demultiplexing, paired-end reads from each sample were were individually mapped to the mm9 genome using BWA-MEM^80^ and converted to bam format and sorted using Samtools^69^. Accessible regions were determined using MACS2 with option -g mm–nomodel–nolambda–broad. Differential peak calling was performed using DESeq2 R pachakge with fdr less than 0.05. Tag counts were normalized to 10 million tags and calculated using annotatePeaks function of HOMER. Unique and overlapping peaks were determined using mergePeaks function of HOMER. ATACseq bigWig tracks were converted from alignments using deepTools^81^ and viewed using IGV. For motif enrichment analysis, genomic coordinates were supplied in BED file format to the HOMER software package^82^, using the “findMotifsGenome.pl” program and default settings.

### Cut and Run analysis

Cut and Run assay was performed as described^56^ with some modifications. 5×10^5^-1×10^6^ activated Treg cells were harvested, cross-linked with 0.5% formaldehyde for 5 min at room temperature, quenched with 0.125 M Glycine for 5 min, and washed twice with cold 1X PBS. Cell pellets were permeabilized in NE1 buffer (0.5mM Spermidine, 20mM HEPES pH7.5, 10mM KCl, 0.1% Triton X, 20% Glycerol, and protease inhibitor) on ice for 10 min, and pelleted by centrifuging at 1,500 rpm at 4°C for 5 min. Cell pellets were resuspended in NE1 buffer. 10 ul of ConA-coated magnetic beads per sample (Bangs Laboratories BP531) were activated by washing twice with Binding Buffer (20mM HEPES pH7.5, 10mM KCl, 1mM CaCl, 1M MnCl2), and then resuspended in Binding Buffer. The beads were then added to permeabilized cell pellets and rotated at room temperature for 10 min. The bead-bound cells were collected using magnet stand, resuspended in CutRun Buffer 1 (20mM HEPES pH7.5, 150mM NaCl, 2mM EDTA, 0.5mM Spermidine, 1mg/ml BSA, and protease inhibitor), and incubated on ice for 5 min. The bead-bound cells were washed once with CutRun Buffer 2 (20mM HEPES pH7.5, 150mM NaCl, 0.5mM Spermidine, 1mg/ml BSA, and protease inhibitor) and resuspended in CutRun Buffer 2. 5ug of antibodies (rabbit anti-mouse Stat3, Millipore 06-596; and rabbit anti-mouse Stat5, R&D AF2168) were added to the bead-bound cells and rotated at 4°C overnight. The immunocomplexes were washed three times with CutRun Buffer 2, and resuspended in CutRun Buffer 2. 700 ng/mL of Protein A/G-MNase (plasmid Addgene ID:123461) were added to the immunocomplexes, mixed, and rotated at 4°C for 1 h. The complexes were washed three times with CutRun Buffer 2, resuspended in CutRun Buffer 2, and incubated on ice to chill down to 0°C. 100mM CaCl2 was added and incubated at 0°C for 30 min to activate MNase. 2X Stop Buffer (340mM NaCl, 20mM EDTA, 4mM EGTA, 0.05% Digitonin, 100 ug/ml RNase A, and 50ug/ml Glycogen) was added, mixed, and incubated at 37°C for 30 min to release the DNA fragments. The mixture was placed on the magnet stand, and supernatant containing digested chromatin was collected. Immunoprecipiated DNA was purified by phenol chloroform extraction. 0.1% SDS and 20mg/ml proteinase K were added to each sample and incubated at 50°C for 1 h, then 65°C for 1 h to reverse cross-linking. An equal volume of phenol chloroform was added, mixed, transferred to Qiagen MaXtract phase-lock tubes, and centrifuged for 5 min at room temperature at 14,000 rpm. An equal volume of chloroform was added and centrifuged for 5 min at room temperature at 14,000 rpm. The top liquid phase was collected, and 2mg/mL glycogen and 100% ethanol were added, mixed, and centrifuged for 10 min at 4°C at 14,000 rpm. DNA pellets were washed once with 100% ethanol, air dried, resuspended in 30 µl of water, and quantified using a Qubit instrument (Invitrogen). Immunoprecipiated DNA was used to prepare libraries using NEBNext Ultra II DNA library prep kit (NEB E7645S). Libraries were sequenced with paired-end 50 on an Illumina Novaseq.

The Cut and Run sequencing data was processed by in-house shell scripts. Briefly, TrimGalore was used to remove adaptor sequence reads. BWA-MEM^80^ was used to align reads to the reference genome mm9. MACS2 was applied to call the binding peaks. P values of differentially enriched binding peaks between different samples were calculated using DESeq2 fdr less than 0.05. Tag counts were normalized to 10 million tags and calculated via the function annotatePeaks of HOMER^82^. Unique and overlapping peaks were determined using mergePeaks function of HOMER. Cut and Run bigWig tracks were converted by deepTools^81^ and viewed using IGV.

### Single cell RNA-Seq analysis

Up to two million cells from each sample were stained with 2µl Totalseq A hashtag antibodies (Biolegend, see supplementary table for details) on ice for 20 minutes, and then cells were washed 3x with washing buffer (PBS, without Ca++ or Mg++, with 0.04% BSA). After staining, viable cells were sorted, and then an equal number of sorted cells from 3 samples (each stained by different hashtag antibodies) were mixed. The cell suspension, 10x barcoded gel beads, and oil were loaded into 10x Chromium™ Single Cell Chip G to capture single cells in nanoliter-scale oil droplets by 10xChromium™ Controller and generate Gel Bead-In-Emulsions (GEMs) (10X Genomics PN-1000127). Single cells were lysed in droplets and single strand cDNA libraries from each single cell were then synthesized by incubation of the GEMs in a thermocycler machine. The reverse transcriptase used nonspecifically added poly C at the 3’ end of the cDNAs and these poly C bound to poly G in “template switch oligos” that were spiked in the solution as a new template. cDNAs were then continuously lengthened with the known 3’ end sequence. After, GEMs containing single cell barcoded-cDNA libraries were pooled together. Pooled cDNAs were cleaned up using DynaBeads MyOne™ Silane beads (Invitrogen 37002D) and were then preamplified by PCR. Preamplified cDNA and hashtag oligos were double selected by SPRIselect beads (Beckman Coulter, PN B23318) to purify cDNA. Next, the remaining supernatant was further selected 2x to purify hashing tag oligos. Gene expression libraries were constructed by PCR amplifying the double-selected preamplified cDNAs with index primers. The hashtag libraries were constructed by PCR amplifying the double-selected preamplified hashtag oligos with index primers. The final constructed 3’-biased gene expression and hashtag libraries were sequenced on an Illumina Novaseq.

For cDNAs derived from mRNAs, cellranger (v.3.0.2,10x Genomics) was used to perform barcode processing and transcript counting after alignment to the mm10 reference genome with default parameters. For cDNAs derived from ADTs, the raw fastq files were reformatted the same way as cDNAs from RNA. The ADT UMI numbers for each antibody in each sample were counted using default settings of CITE-seq-Count (v.1.4.2). The single-cell transcriptome data and single-cell CITE-seq data were integrated via Weighted Nearest Neighbors (WNN) approach in the Seurat (v.4.0.0) R package^83^. The cells that expressed fewer than 200 genes were filtered out and all genes that were not detected in at least three single cells were excluded. The processed data was normalized using Seurat’s ‘NormalizeData’ function, which used a global scaling normalization method, LogNormalize, to normalize the gene expression measurements for each cell to the total gene expression. Highly variable genes were then identified using the function ‘FindVariableGenes’ in Seurat. The anchors were identified using the ‘FindIntegrationAnchors’ function, and the matrices from different samples were integrated with the ‘IntegrateData’ function. The variation arising from library size and percentage of mitochondrial genes was regressed out using the function ‘ScaleData’ in Seurat. Principal component (PC) analysis was performed using the Seurat function ‘RunPCA’, and the K-nearest neighbor graph was constructed using the ‘FindNeighbors’ function in Seurat with the number of significant PCs identified from PCA analysis. Clusters were identified using the ‘FindClusters’ function with a resolution of 0.8. The clusters were visualized in two dimensions with UMAP. The normalization, integration, and clustering were performed under standard Seurat workflow. Differential gene expression analyses were carried out using the Seurat function ‘FindMarkers’. Briefly, we performed the Wilcoxon rank-sum test with the default threshold of 0.25 for log2 fold change and a filter for the minimum percent of cells in a cluster greater than 25%. Differentially expressed genes were isolated by comparing significantly upregulated genes and downregulated genes defined as adjusted P-value, Padj < 0.05. Top differentially expressed genes were visualized by heatmap via ComplexHeatmap (v2.11.1) package in R. Expression of target genes were represented as violin plots by function ‘VlnPlot’, dot plots by function ‘DotPlot’, or UMAPs by function ‘FeaturePlot’ in Seurat.

### ChIP-Seq analysis

The sequencing raw reads of GSE113054^54^ were downloaded from the GEO database using the SRA Toolkit (v.2.11.2). Raw single-end sequencing data were mapped to mm9 genome with BWA-MEM^80^. ChIP-seq peaks were called using MACS2^84^ with p value cut off at 1×10^−5^. The total input DNA was used as the control for peak-calling. Differential peak calling was performed using DESeq2 with FDR less than 0.05. Tag counts were normalized to 10 million tags and calculated using annotatePeaks function of HOMER^82^. Unique and overlapping peaks were determined using mergePeaks function of HOMER. ChIPseq bigWig tracks were converted from alignments using deepTools^81^ and viewed using IGV.

### Real time PCR

Total RNA was isolated using miRNeasy Micro Kits according to the manufacturer’s instructions (QIAGEN 217084). cDNA was synthesized with the iScript Reverse Transcription Supermix (Bio-Rad 170-8841), and real-time PCR was performed on a Bio-Rad CFX Connect Real-Time System using SsoAdvanced Universal SYBR Supermix (Bio-Rad 172-5275). Reactions were run in duplicate. Gene-of-interest (GOI) CTs were normalized to β2M CTs.

### Crispr knock down

CRISPR-Cas9 RNP nucleoporation was carried with adaptations^85,86^. Briefly, three unique guides specific to each gene were designed using IDT’s online design tool specifically targeting the 5’ end of each target. Alt-R tracrRNA and guides were purchased from IDT and dissolved in Nuclease-Free Duplex Buffer (IDT 11-05-01-03) at 200 μM. Individual crRNAs and tracrRNA were mixed at equimolar concentrations in sterile PCR tubes and annealed by heating to 95°C in a thermocycler for 5 minutes followed by cooling to room temperature. To generate RNP complexes, 1 μL of each of the three unique sgRNAs targeting each gene were combined, added to 2 μL of TruCut Cas9 Protein v2 (5 ng/μL, Thermo Fisher Scientific A36498), and incubated for 10 minutes at room temperature. Negatively selected mouse naïve CD4^+^ T cells, were incubated for 24 hours in RPMI-10 supplemented with IL-7 (5 ng/mL, Peprotech 217-17). T cells were then rinsed with PBS and 10 × 10^6^ cells were resuspended in 20 µl of P4 primary cell nucleofection solution (P4 Primary Cell 4D-Nucleofector X Kit S, Lonza V4XP-4032). 20 μL of resuspended T cells were then gently mixed with 5 µL of RNP complex and incubated for 2 minutes at room temperature. The volume of each T cell/RNP mix was electroporated in a Lonza 4D Nucleofector 16-well cuvette and received pulse DS137. Post-nucleofection, pre-warmed IL-7-containing media was added directly to the cuvettes and cells were gently resuspended and aliquoted into 24 well plates containing pre-warmed IL-7-containing media at a density of 1-2 million cells/mL and kept in culture for 5 days at 37 °C in a humidified incubator with 5% CO_2_. sgRNA species used are listed in Supplementary Table.

### Th17 transfer colitis and histology

Th17 cells were polarized as described. Live GFP(IL-17A)^+^ cells were sorted and a total of 4 x 10^5^ cells were injected intraperitoneally into age-matched *Rag1*^-/-^ recipients. iTreg cells were polarized as described; 3 weeks after the Th17 transfer, GFP(Foxp3)^+^ cells were sorted and 1 x 10^6^ cells were transferred into recipient mice intraperitoneally. Mice were monitored regularly for signs of disease and were weighed weekly. 6 weeks after Th17 transfer, mice that did not receive iTreg cells were sacrificed according to our IACUC-approved protocol (average weight below 80%). Mice that received iTreg cells survived until resolution of disease at 12 weeks. At both time points, colons were recovered and processed as described. Samples were scored by a pathologist in a blinded fashion as previously described (Maynard)^16^.

### Statistics

Statistics were performed using Prism (GraphPad). Data are expressed as mean ± standard error of the mean. p values were calculated using two-tailed Student’s t tests and one-way or two-way ANOVA tests with Tukey’s post hoc multiple comparisons analysis. A p value of <0.05 was considered significant. See figure legends for details.

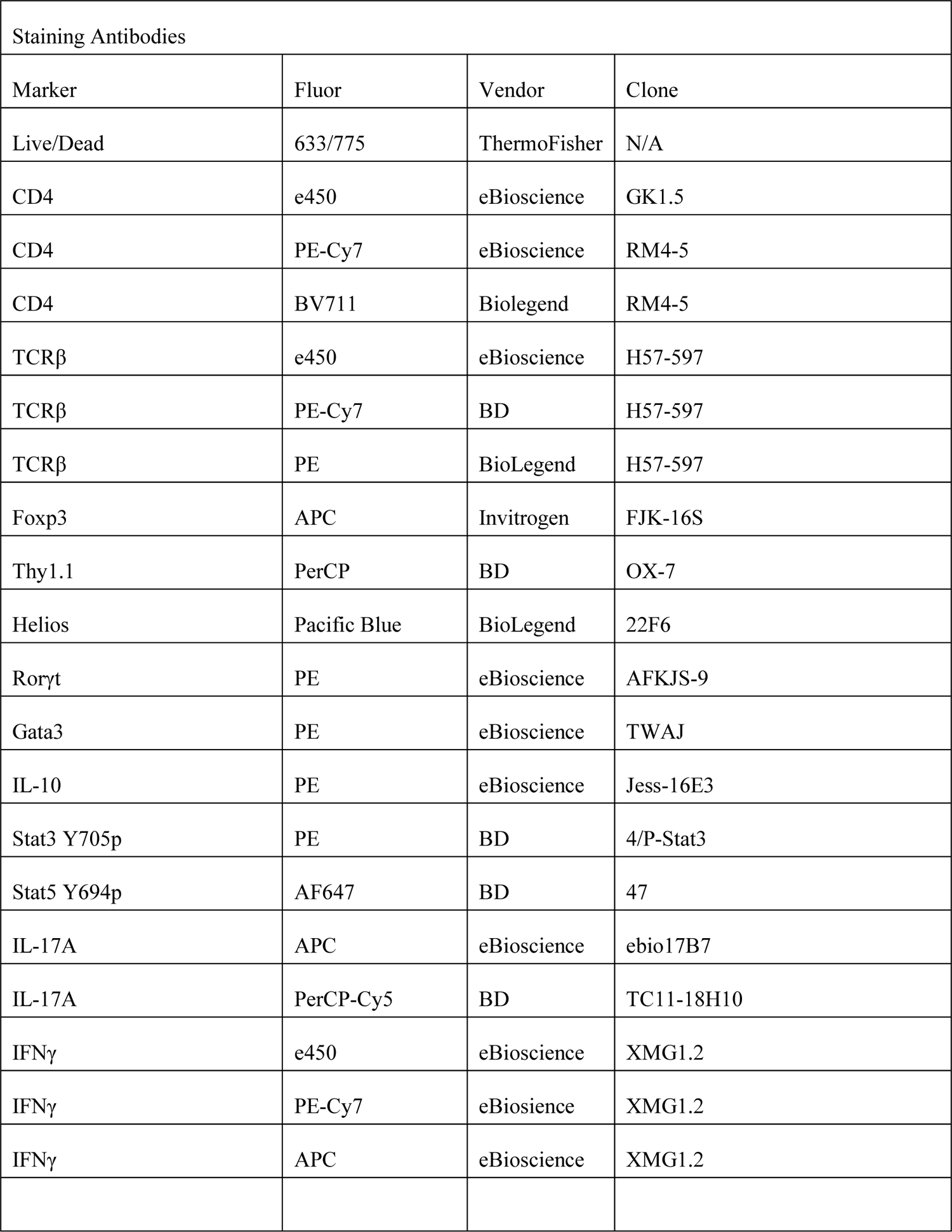
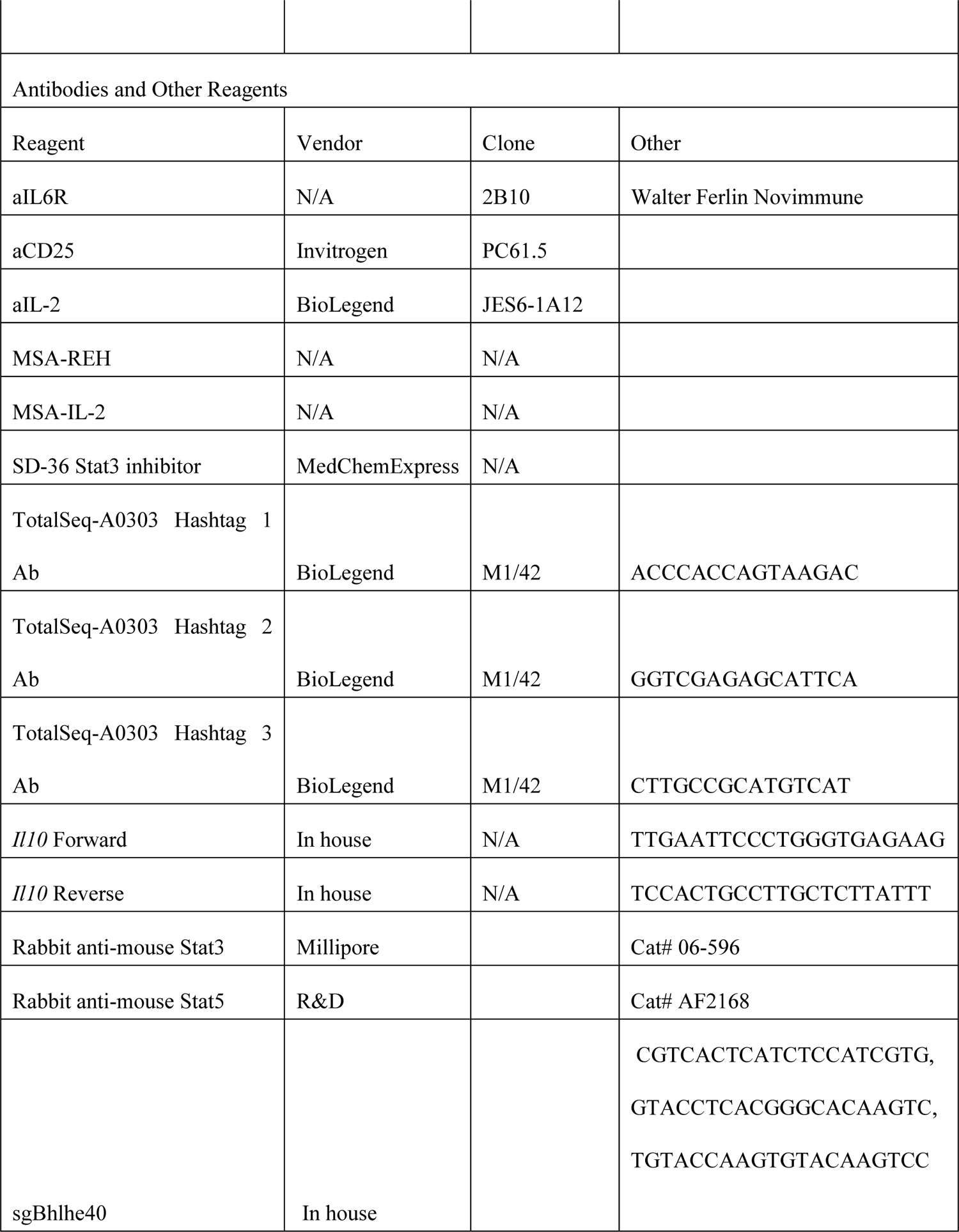

## Supplemental figure legends

**Fig. S1.**
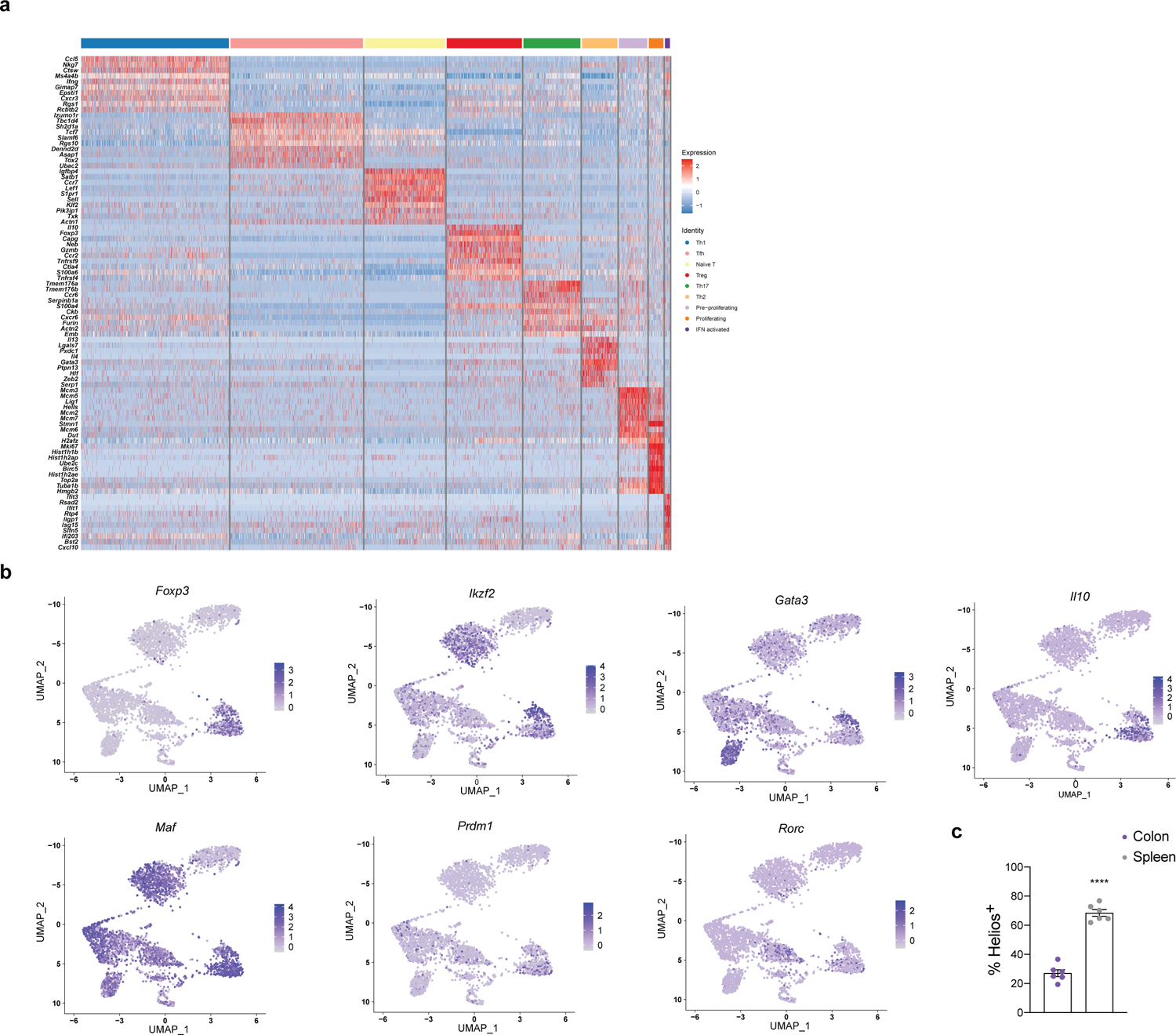
CD4^+^ T cell single cell profiling reveals expression of factors in distinct sub-populations. **a-b,** Single cell RNA-seq was performed on live CD4^+^TCRβ^+^ T cells isolated from the colonic lamina propria of naïve C57BL/6 mice. Heatmap (**a**) shows cluster-defining genes. UMAP overlays (**b**) show expression of indicated genes. Data is representative of two biologically independent experiments. **c,** Quantification of frequency of Helios^+^ Treg cells (gated on CD4^+^TCRβ^+^*Foxp3*GFP^+^) isolated from the colonic lamina propria (purple) or spleens (grey) of 10BiT.*Foxp3*GFP mice (n=6 for both colon and spleen, mean ± SEM; ****P<0.0001). Statistical differences were tested using unpaired Student’s t-test (two-tailed).

**Fig. S2.**
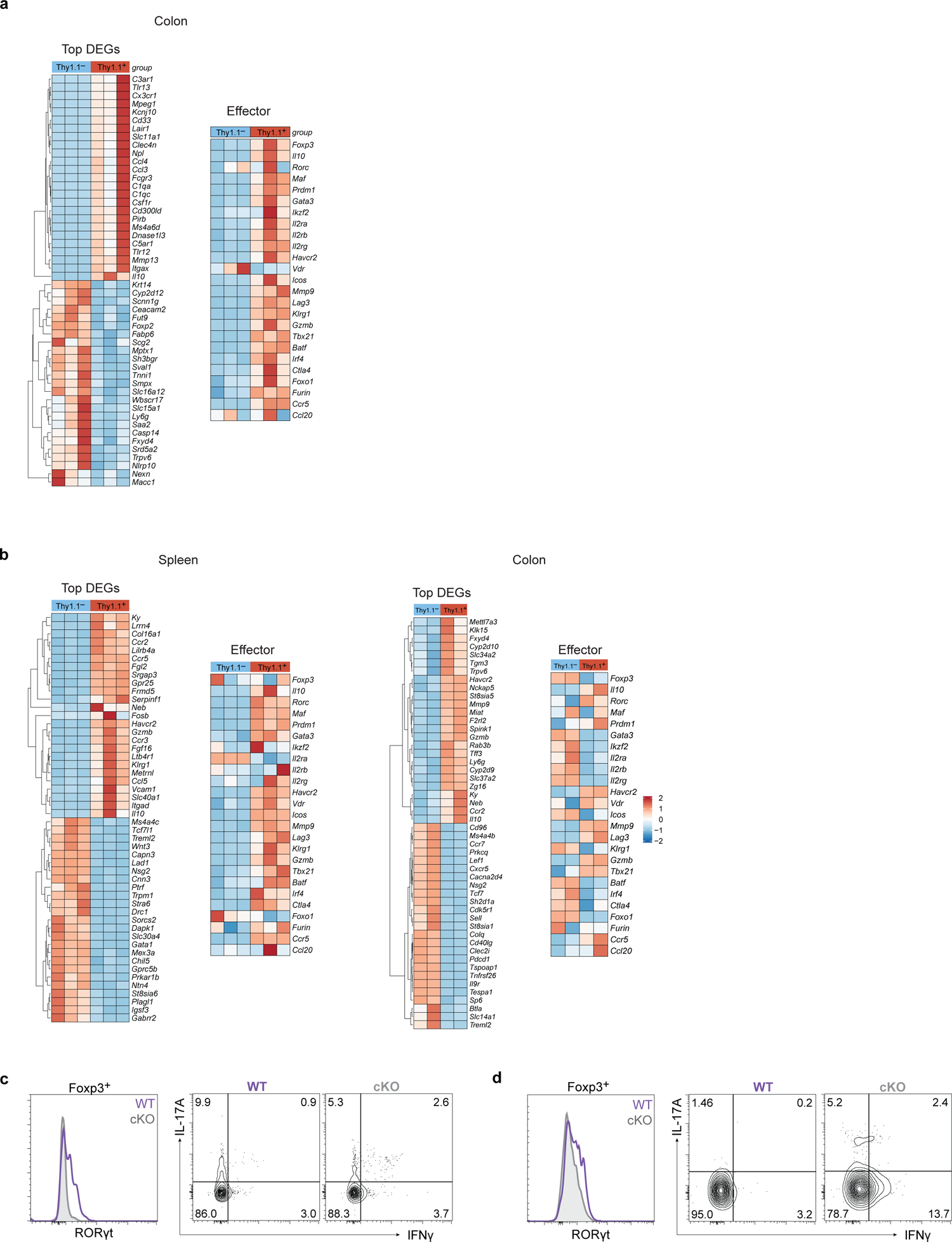
Stat3 signaling and Rorγt expression are enriched in colonic Treg cells. **a**, Treg cells (gated on CD4^+^TCRβ^+^*Foxp3-*GFP^+^Thy1.1^-^ and CD4^+^TCRβ^+^*Foxp3-* GFP^+^Thy1.1^+^) were sorted from colonic lamina propria of 10BiT.*Foxp3-*GFP mice and subjected to RNA-sequencing. Heatmaps (larger) showing relative expression (z-score) of differentially expressed genes (FDR<0.1) between Thy1.1^-^ and Thy1.1^+^ Treg cells in the colon (n=3). Heatmaps (smaller) showing relative expression (z-score) of select effector-associated transcripts. **b**, Treg cells (gated on CD4^+^TCRβ^+^*Foxp3-*GFP^+^Thy1.1^-^ and CD4^+^TCRβ^+^*Foxp3-* GFP^+^Thy1.1^+^) were sorted from colonic lamina propria (right) and spleen (left) of 10BiT.Foxp3eGFP mice and subjected to RNA-sequencing. Heatmaps (larger) showing relative expression (z-score) of differentially expressed genes (FDR<0.1) between Thy1.1^-^ and Thy1.1^+^ Treg cells in the colon (n=2) and spleen (n=2). Heatmaps (smaller) showing relative expression (z-score) of select effector-associated transcripts. **c-d,** Histograms (left) showing Rorγt expression in colonic Treg cells (gated on CD4^+^TCRβ^+^Foxp3^+^) from *Stat3*^CD4Cre^ (WT, purple) and *Stat3*^ΔCD4Cre^ (cKO, grey) mice (**c**) or *Stat3*^Foxp3Cre^ (WT, purple) and *Stat3*^ΔFoxp3Cre^ (cKO, grey) mice (**d**). Flow cytometric profiles (**b, c**) showing IL-17A and IFNγ expression in colonic CD4^+^TCRβ^+^ T cells from WT and cKO mice. Histograms and profiles are representative of three independent experiments.

**Fig. S3.**
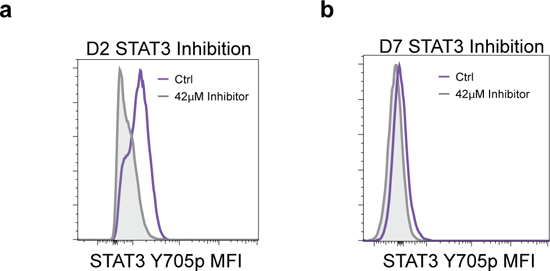
Stat3 inhibition by SD-36 effectively reduces Stat3 Y705p. **a-b**, Naïve CD4 T cells were isolated from 10BiT.*Foxp3*GFP mice and polarized in iTreg conditions as in Fig. 3a. but 42µM SD-36 (Stat3 inhibitor) or DMSO (control, left) was added on each day to independent wells. Histograms show Stat3 Y705p on day 8 when 42µM of inhibitor (grey) or DMSO (purple) was added on day 2 (**a**) or day 7 (**b**) of polarization. Experiment performed twice.

**Fig. S4.**
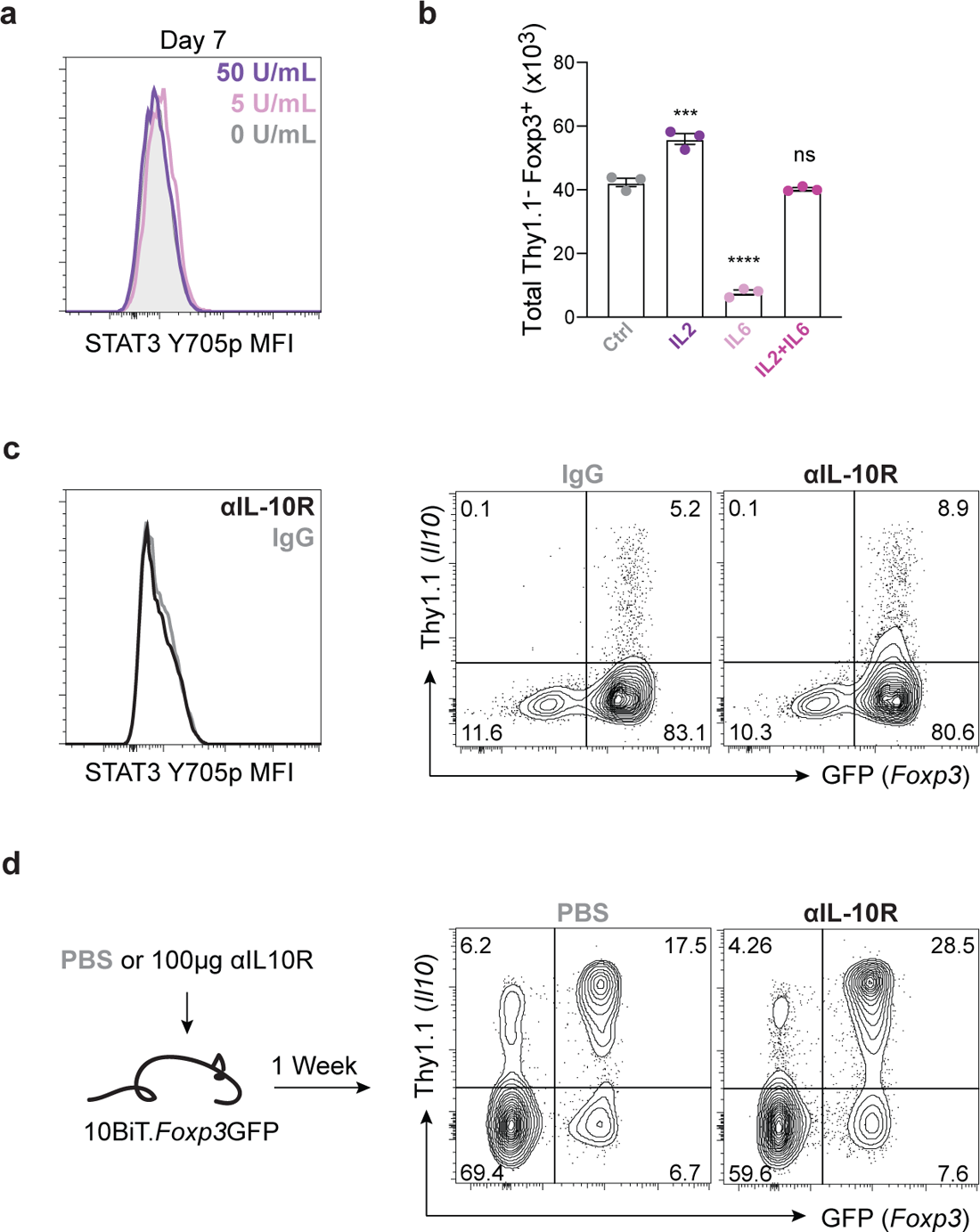
Early Treg Stat3 activation is controlled by IL-2 exposure. **a**, Naïve CD4 T cells were isolated from 10BiT.*Foxp3*GFP mice and polarized in vitro in iTreg conditions using 0 (grey), 5 (pink) or 50 (purple) U/mL IL-2 during the induction phase (left) and 50 U/mL during re-stimulation. Histogram shows Stat3 Y705p on day 7 for each condition (gated on live CD4^+^*Foxp3*GFP^+^). **b,** Naïve CD4 T cells were isolated from 10BiT.*Foxp3*GFP mice and polarized *in vitro* in iTreg conditions with 50U/mL IL-2 (Control, grey), 50U/mL IL-2 + 10µg/mL αIL-6Rα (IL-2 + anti-IL-6a, purple), 10µg/mL αCD25 nd 0.1ng/mL IL-6 (anti-IL-2 + IL-6, pink), or 50U/mL IL-2 + 0.1ng/mL IL-6 (IL-2 +IL-6, magenta). Quantification shows number of Thy1.1^-^ *Foxp3*GFP^+^ cells on day 8 (n=3, mean ± SEM; ns (nonsignificant), ***P<0.001, ****P<0.0001). **c,** Naïve CD4 T cells were isolated from 10BiT.*Foxp3*GFP mice and polarized *in vitro* in iTreg conditions with 10µg/mL αIL-10R (black) or IgG control (grey). Histogram shows Stat3 Y705p on day 2. Flow cytometric profiles show Thy1.1 versus *Foxp3*GFP expression on day 8. Experiment performed twice. **c,** 10BiT.*Foxp3*-GFP mice were treated with 100µg αIL-10R (black) or IgG (grey) on day 0. On day 7, animals were sacrificed and lymphocytes were collected from the colonic lamina propria. Flow cytometric profiles show Thy1.1 versus *Foxp3*GFP expression (gated on CD4^+^TCRβ^+^). Statistical differences were tested using one-way ANOVA.

**Fig. S5.**
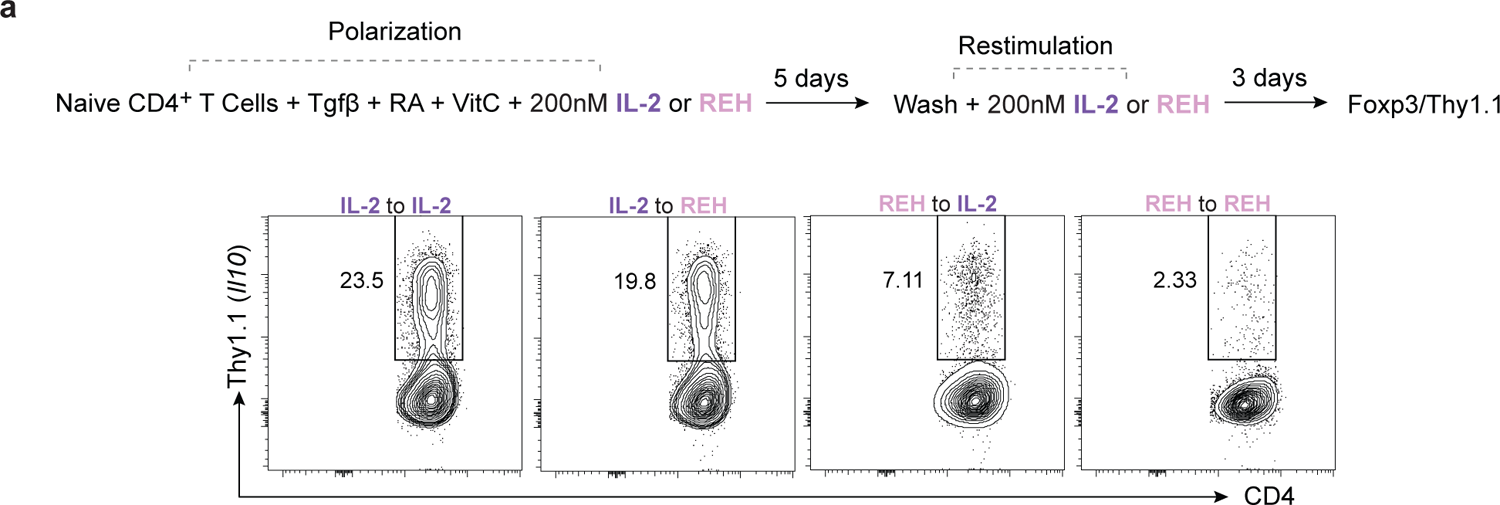
The effects of REH signaling on Treg cells are not dependent on Stat3. **a**, iTreg cells were polarized with either 200nM MSA-IL-2 (purple) or 200nM MSA-REH (pink) during the induction phase. On day 5, cells were restimulated with either 200nM MSA-IL-2 (purple) or MSA-REH (pink) for an additional three days. On day 8, cells were harvested and assessed by flow cytometry. Flow cytometric profiles show expression of Thy1.1 and CD4 (gated on CD4^+^*Foxp3*GFP^+^). Experiment performed twice.

**Fig. S6.**
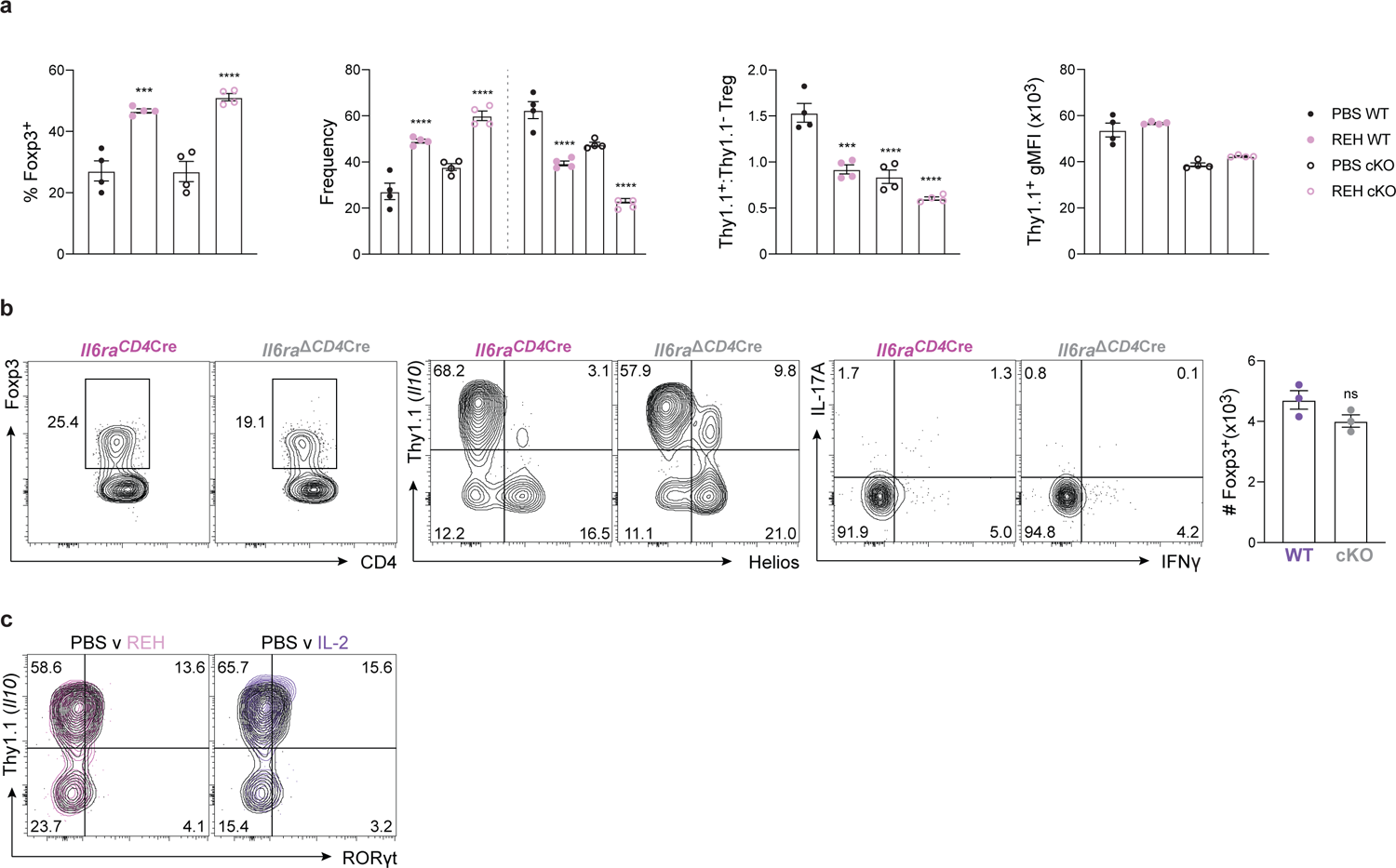
IL-2 receptor signaling specifically supports Rorγt iTreg cells in the colon. **a**, Gender matched *Stat3*^Foxp3Cre^ (WT, closed circles) and *Stat3*^ΔFoxp3Cre^ (cKO, open circles) littermate mice were treated I.P. with 30µg/mL of REH (pink) or PBS (black) on days 0, 3, and 6. Animals were sacrificed on day 7 and cells were isolated from the colonic lamina propria. Quantifications show frequency of *Foxp3*YFP^+^CD4^+^ T cells (left, gated on CD4^+^TCRβ^+^), Helios^+^ and Helios^-^ Treg cells (center left, gated on CD4^+^TCRβ^+^Foxp3^+^), the ratio of Thy1.1^+^ to Thy1.1^-^ Treg cells (center right, gated on CD4^+^TCRβ^+^Foxp3^+^), and Thy1.1+ gMFI (right, gated on CD4^+^TCRβ^+^Foxp3^+^Thy1.1^+^) (n=4, mean ± SEM; ***P<0.001, ****P<0.0001). Statistical differences were tested using one- or two-way ANOVA. **b,** Colonic lamina propria lymphocytes were isolated from gender matched *Il6raCD4*Cre^-^ (WT, purple) and *Il6raCD4*Cre^+^ (cKO, grey) littermate mice. Flow cytometric profiles show expression of Foxp3 versus CD4 (left, gated on CD4^+^TCRβ^+^), Thy1.1 versus Helios (middle, gated on CD4^+^TCRβ^+^*Foxp3*YFP^+^), and Il-17A vs Ifnγ (right, gated on CD4^+^TCRβ^+^). Quantification depicts the number of colonic Foxp3^+^ Treg cells in each group (n=3, mean ± SEM; ns (nonsignificant). **c,** 10BiT mice were treated with PBS (black), 5µg of MSA-IL-2 (purple), or 5µg MSA-REH (pink) on days 0, 2, 4, and 6 and sacrificed on day 7. Lymphocytes were isolated from the colonic lamina propria and assessed by flow cytometry. Flow cytometric profiles show overlays of PBS and REH treat animals (left) or PBS and IL-2 treated animals (right). Experiment performed twice. Statistical differences were tested using unpaired Student’s t-test (two-tailed).

**Fig. S7.**
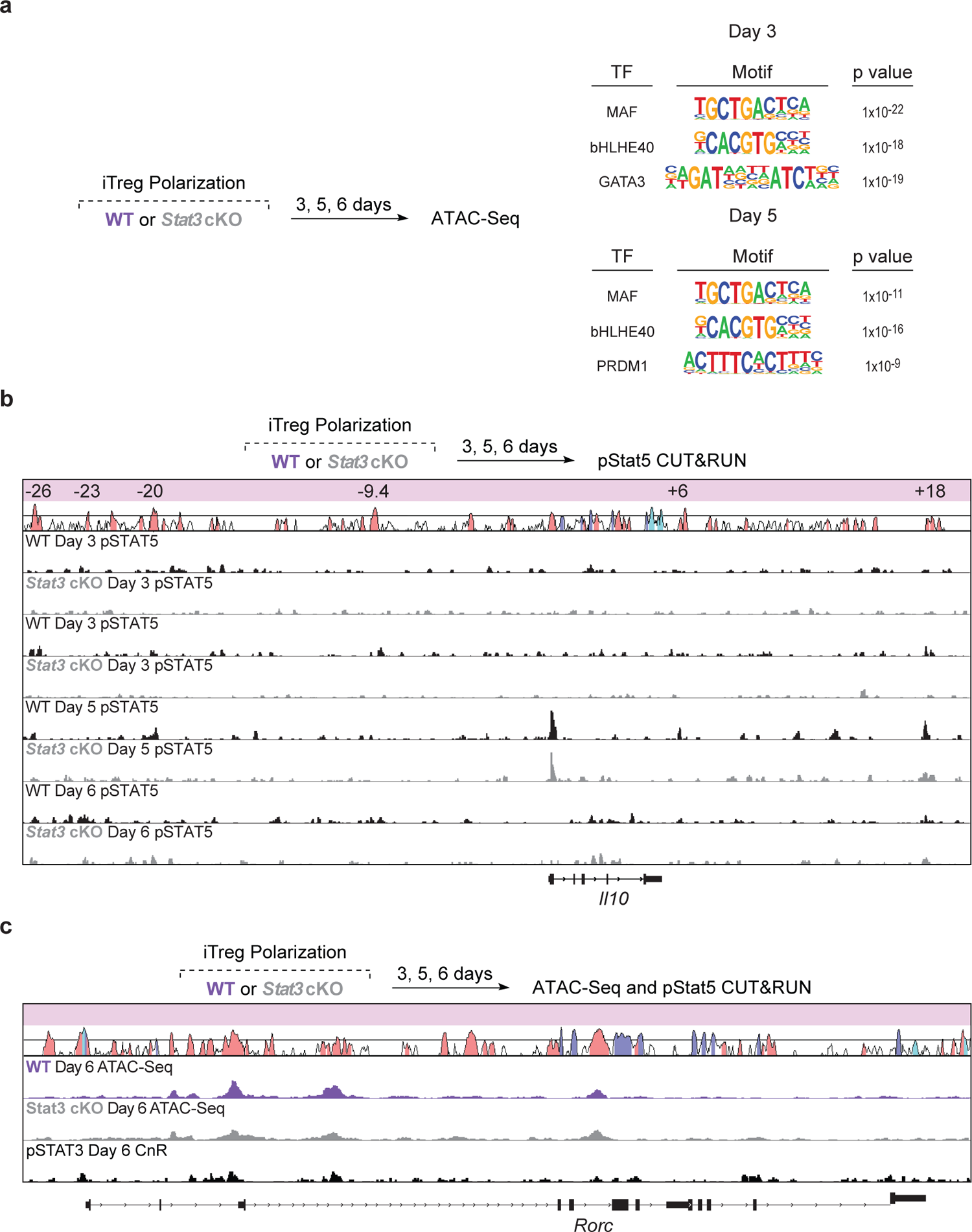
Stat3 and Stat5 control effector loci. **a**, Naïve CD4^+^ T cells were isolated from *Stat3*^CD4Cre^ (WT, purple) or *Stat3*^ΔCD4Cre^ (cKO, grey) and polarized as in Fig. 3a with 50U/mL IL-2. ATAC-seq was then performed on live cells on days 3, 5, and 6. Tables showing a selection of enriched motifs, their associated transcription factor, and p value for each timepoint shown. **b**, Naïve CD4^+^ T cells were isolated from *Stat3*^CD4Cre^ (WT, black) or *Stat3*^ΔCD4Cre^ (cKO, gray) and polarized as in Fig. 3a with 50U/mL IL-2. On days 3, 5, and 6, cells were harvested and subjected to pStat5 CUT&RUN. Pink asterisks show areas of enriched pStat5 binding in WT cells at *Il10* (shown below each day’s tracks). **c**, Naïve CD4^+^ T cells were isolated from *Stat3*^CD4Cre^ (WT, purple) or *Stat3*^ΔCD4Cre^ (cKO, grey) and polarized as in Fig. 3a with 50U/mL IL-2. ATAC-seq was then performed on live cells on days 3, 5, and 6 (only day 6 shown here). Naïve CD4^+^ T cells were also isolated from C57BL/6 WT mice (black) and polarized as in Fig. 3a with 50U/mL IL-2. pStat3 CUT&RUN was then performed on days 3, 5, and 6 (only day 6 shown here). Pink asterisks denote DA between Stat3 WT and cKO iTreg cells (ATAC-Seq) or pStat3 binding regions (CnR) at *Rorc*. Data represent the combined analysis of two biologically independent samples.

